# Unlocking Ultrasensitive Phosphoproteomics of Nanoscale Samples: A Rapid IMAC Enrichment Platform Using Ti-PAN Tip

**DOI:** 10.64898/2026.01.08.698355

**Authors:** Ci Wu, Yao Lin, Xingyao Wang, Jiawei Gong, Bing Liu, Yuting Lu, Feitai Tang, Mei Yang, Ge Lin, Jue Zhang, Shen Zhang

**Author notes:** These authors contributed equally to this work (Ci Wu, Yao Lin, Xingyao Wang, Jiawei Gong).

## Abstract

Protein phosphorylation is a critical mechanism in cellular signal transduction, playing essential roles in various cellular processes. However, significant challenges of their low abundance, high dynamic range, and complexity are especially prominent for accurate phosphoproteome analysis in nanoscale samples, such as oocytes, where misregulation of phosphorylation can lead to infertility. In this study, we developed a novel phosphopeptide enrichment tip (Ti-PAN) using highly hydrophilic textile PAN fibers, enabling rapid in-situ one-tip IMAC enrichment in just 8 minutes. This approach not only minimizes sample loss but also enhances analytical sensitivity and is easily adaptable to automation for high-throughput applications, demonstrating excellent recovery (>94%) and outstanding selectivity for phosphopeptides with remarkable resistance to interference up to 1:5000. By integrating it with a compatible sample preparation workflow based on n-dodecyl β-D-maltoside (DDM) lysis with the high-sensitivity MS data acquisition method of turboDDA, comprehensive mapping of phosphosites in microsamples was achieved, including those from the ERK interactome, mouse brain slices, and oocytes. Using this approach, we identified 2,139 to 11,296 phosphosites across 645 to 2,168 phosphoproteins from samples ranging from 100 to 100,000 cells. Furthermore, we successfully applied this strategy to analyze the phosphoproteome of mouse oocytes at different developmental stages, using as few as 5 oocytes per sample. This led to the identification of 2,709 phosphosites on 701 phosphoproteins, revealing several key kinases involved in meiotic resumption and oocyte maturation, including STK4 (serine/threonine kinase 4) and DYRK3 (dual specificity tyrosine phosphorylation-regulated kinase 3). To our knowledge, this represents the deepest coverage of phosphoproteomes from such minimal oocyte material reported to date, offering valuable mechanistic insights into developmental controls during oocyte maturation. This Ti-PAN-based method provides a robust, versatile, and automatable platform for large-scale, highly sensitive phosphopeptide enrichment, substantially advancing phosphoproteomic studies in complex biological systems.

## Introduction

Protein phosphorylation, as one of the most significant and extensively studied post-translational modifications, playing a central role in a wide range of physiological and pathological processes, including cell signaling, metabolic regulation, DNA repair, and apoptosis.^[1–3]^ Phosphoproteomics has emerged as an essential tool for exploring cellular functions and disease mechanisms, showcasing considerable potential in the investigation of complex diseases such as cancer and neurodegenerative diseases, genetics and developmental disorders.^[4–8]^ However, the low abundance, high dynamic range, and complexity of phosphorylation in biological samples present significant challenges for accurate identification and comprehensive phosphoproteome analysis. These issues are especially prominent when analyzing precious microscale samples, such as tissue sections from clinical specimens, affinity purification-mass spectrometry (AP-MS) sample, early embryos, and circulating tumor cells.^[9–10]^. In these cases, sample loss and the low abundance of phosphorylated peptides are highly vulnerable to matrix effects and technical limitations, often leading to incomplete or inaccurate analytical results. Moreover, due to cellular heterogeneity, achieving high-throughput and in-depth phosphoproteomic analysis in microscale samples such as single cells is another key factor. For instance, dynamic phosphorylation modifications during the development of mouse oocytes are essential for understanding oocyte maturation and meiosis, as misregulation of phosphorylation due to kinase mutations can impair oocyte quality and result in female infertility.^[11]^ Currently, due to technical limitations and the scarcity of material, analyzing the temporal phosphorylation dynamics of mammalian oocyte maturation requires large numbers of mouse oocytes (tens of thousands), a challenge that makes similar analysis impossible for human oocytes and embryos. Consequently, developing phosphoproteomic methods with enhanced sensitivity, minimized sample loss, and high throughput has become a central focus in the field of microscale phosphoproteomics.

Currently, methods for enriching phosphorylation primarily include classical techniques such as Immobilized Metal Ion Affinity Chromatography (IMAC)^[12–13]^ and TiO₂ beads^[14–16]^. However, traditional IMAC faces limitations such as low chelation site density and high hydrophobicity of the matrix, susceptibility to nonspecific adsorption, and relatively low recovery rates. TiO₂-based methods often suffer from limited selectivity and throughput, and the multiple centrifugation and transfer steps involved may lead to significant sample loss, an issue that is especially critical when handling microscale samples. Recently, by integrating enrichment materials with several innovative sample preparation strategies and rapid proteomic workflows, such as techniques involving microwaves^[17]^ and ultrasound^[18]^, have developed to accelerate the digestion process and effectively address the challenges in microscale phosphoproteomics^[19]^. Kelly et al.^[20]^ employed an optimized Suspension trapping (S-Trap) approach, where trifluoroacetic acid is substituted for phosphoric acid, providing a simple and effective method to prepare low-abundance, membrane-rich microscale samples for phosphoproteomics.

To further enhance phosphoproteome identification, researchers have also incorporated advanced mass spectrometry (MS) acquisition methods, such as data-independent acquisition (DIA).^[21]^ Chen et al.^[22]^ developed a simple and rapid one-pot phosphoproteomics workflow utilizing TiO₂ bead enrichment in a single-tube format, integrated with DIA-MS for microscale phosphoproteomic analysis. Although this approach enabled scalable phosphopeptide analysis from as few as 2,500 cells, it depends on an existing phosphorylation database library. Recently, a data-dependent acquisition method combining fast MS/MS scanning and no dynamic exclusion (turboDDA) has been developed in our group, which demonstrates greater sensitivity and accuracy compared to traditional data-dependent acquisition with dynamic exclusion (DEDDA) and DIA, particularly for microscale samples.^[23, 30]^

Overall, ongoing innovations in enrichment strategies and detection technologies offer promising prospects for microscale phosphoproteomic analysis. However, challenges such as low enrichment efficiency, matrix interference from current enrichment materials, and sample loss during preparation continue to impede further progress. The development of highly selective enrichment materials and enhancements in detection sensitivity are essential to advance both fundamental research and clinical applications of phosphoproteomics.

In this study, we developed a rapid in-situ one-tip IMAC phosphoproteomics enrichment workflow, integrated with turboDDA-MS, for nanoscale phosphoproteomic analysis. The enrichment material Ti-PAN was fabricated from textile PAN fiber, which is highly hydrophilic and readily functionalized with phosphate groups covalently. Through systematic optimization of nanosample preparation and enrichment conditions, we achieved in-depth identification of phosphosites from low-input amounts of 100 to 100,000 HeLa cells. Furthermore, our strategy was successfully applied to phosphoproteomic analysis of nanoscale samples, including AP-MS samples and mouse brain slices. Notably, we successfully analyzed the phosphoproteome of mouse oocytes at different maturation stages using as few as five oocytes per sample, showcasing the robust nanosample phosphopeptide enrichment capabilities of our material. This unprecedented level of sensitivity makes it possible to uncover novel stage-specific phosphorylation sites and associated kinases, thereby paving the way for reliable phosphoproteomic analysis of even more scarce and valuable clinical samples such as single human oocytes and embryos in the future.

## Experimental Section

### Preparation of Ti-PAN Material

Polyacrylonitrile fibers (PAN) was soaked in a 100 mM 1,3-diaminopropane solution and heated at 60 ℃ for 8 h. After the reaction, the fiber was thoroughly washed with H_2_O to remove excess 1,3-diaminopropane and dried for subsequent functionalization. Next, 10 mg of the obtained fiber was added to 10% (v/v) glutaraldehyde in 100 mM PBS (pH 8.0) and stirred at room temperature for 6 h. Subsequently, the fiber was washed with H_2_O. After the extra water was squeezed out, the fiber was incubated in a solution containing 2 mg/mL AMPA, 10 mg/mL NaCNBH₃, and 100 mM PBS (pH 8.0) under stirred at room temperature for 6 h followed by washing with H_2_O. Finally, the fiber was immersed in 100 mM Ti(SO_4_)_2_ aqueous solution and reacted overnight at room temperature to get Ti-PAN. Excess Ti^4+^ were removed by extensive washing with H_2_O.

### Sample Preparation of Real Complex Sample

HeLa cells (serial dilution was performed to obtain different numbers with 100, 1000, 10,000 and 100,000), mouse oocytes and mouse brain slices were lysed with lysis buffer (0.01% DDM, 1% cocktail, 1% phosphatase inhibitor, 100 mM TEAB pH 8.5) and ultrasonicated for 10 cycles (30 s on/30 s off) at 4 ℃. Then, the obtained lysates were heated at 95 ℃ for 5 min and the cellular lysate was reduced with 2 mM DTT for 30 min at 37 ℃ followed by alkylating with 5 mM IAA for 30 min at room temperature in dark. Then, proteins were digested with trypsin at an final concentration of 10 ng/μL, and incubated at 37 ℃ for 4 h. The same amount of trypsin was added again at 37 ℃ overnight.

### Enrichment of Phosphopeptides by Ti-PAN tip

One mg of Ti-PAN material was packed into an empty low retention pipette tip. The Ti-PAN tip was preconditioned by adding 20 μL of loading buffer (60% ACN, 6% TFA, 10 mM lactic acid) to a centrifuge tube followed by aspirating-dispensing buffer for 10 cycles, repeated three times for equilibration. Peptide samples in loading buffer were loaded onto Ti-PAN with 20 aspirate-dispense cycles. Sequential washing was performed using 10 or 20 μL of Wash Buffer I (loading buffer/ACN, v/v 1:1) and Wash Buffer II (0.5% TFA), through 10 cycles each, repeated twice for both buffers. Phosphopeptides were then eluted using 20 μL of elution buffer (50% ACN, 5% NH_3_·H_2_O) with 20 aspirate-dispense cycles for three times into the sample vial insert. Finally, eluates were directly vacuum-dried in situ for LC-MS/MS analysis without transfer steps.

### NanoLC-MS/MS

MS analysis was performed using a Thermo Fisher Orbitrap Eclipse Tribrid mass spectrometer coupled with a FAIMS Pro Interface, employing the turboDDA acquisition method. Enriched phosphopeptides were directly loaded onto a 2 cm PEPMAP trap column on a Vanquish Neo UHPLC system (ThermoFisher Scientific). Separation utilized a 25-cm PepMap analytical column (ThermoFisher Scientific) under two gradient conditions: (i) 102-min linear gradient from 3% to 38% buffer B (80% ACN, 0.1% formic acid), followed by 10-min maintenance at 100% B; or (ii) 51-min gradient (3-38% B) with 5-min equilibration at 100% B. FAIMS compensation voltages cycled through -35 V, -45 V, and -65 V. Full-scan MS1 spectra (350-1500, m/z) were acquired in the Orbitrap (resolution: 60,000; AGC target: standard; max injection time: Auto; RF lens: 50%). Top 100 precursors were selected for MS/MS without dynamic exclusion. Fragment spectra (isolation window: 1.6 m/z) were generated in the linear ion trap via HCD (35% CE) at rapid scan rate (AGC: standard; max IT: Auto; centroid data mode).

DIA analysis was also performed on the same Thermo Fisher Orbitrap Eclipse Tribrid mass spectrometer. The FAIMS interface was set to a compensation voltage of -45 V. Full MS scans were acquired in the Orbitrap with a resolution of 60,000, automatic gain control (AGC) target set to “Standard”, maximum injection time (MaxIT) in “Auto” mode, RF lens at 30%, and a mass range of *m/z* 350-1,250. MS/MS spectra were collected in the Orbitrap with an isolation window of 9 *m/z*, higher-energy collisional dissociation (HCD) activation at 32% collision energy, AGC target of 2000%, MaxIT in “Auto” mode, RF lens at 50%, and centroid data type.

## Results and discussion

### Synthesis and Characterization of Ti-PAN material

In this study, polyacrylonitrile (PAN) was used as the matrix due to its strong hydrophilicity and excellent plasticity, facilitating weak adsorption toward non-phosphopeptides. To achieve efficient, highly sensitive, and low-loss enrichment of phosphorylated peptides, Ti^4+^ ions were grafted onto PAN, and the resulting material was fabricated into pipette tips of different sizes, which facilitates high-throughput automated operations (Figure 1A). Briefly, PAN was first treated with diamines to introduce sufficient −NH_2_ groups for subsequent modification, followed by activation with glutaraldehyde based on a previously reported workflow.^[24]^ A heterofunctional reagent, AMPA, was then introduced to yield functional −PO_4_^3−^ groups, which immobilize Ti^4+^ via chelation, resulting the Ti-PAN material. Finally, Ti-PAN was packed into empty pipette tips for in situ enrichment of phosphopeptides. SEM images of the PAN fibers revealed that their fibrous morphology was largely preserved during the reactions, with the fiber diameter remaining at approximately 10 μm (Figure 1B–1E). In addition, after immobilization of Ti⁴⁺ ions, the surface of the PAN fibers changed noticeably from smooth to rough. Further EDX spectroscopic analysis showed that Ti was evenly distributed throughout the Ti-PAN fibers, demonstrating the successful chelation of Ti⁴⁺ ions to the PAN fibers (Figure 1G). The corresponding elemental composition, including carbon, oxygen, nitrogen, phosphorus, and titanium, is shown in Figure S1. FT-IR spectroscopy was used to confirm each step of the Ti-PAN preparation. As shown in Figure 1F, absorption bands at 1556 cm^−1^ (stretching) and 3297 cm^−1^ (bending) were attributed to −NH_2_ vibrations. The strong band at 1672 cm^−1^ was assigned to C=O vibrations, while the peak at 1033 cm^−1^ corresponded to −PO_4_^3−^ vibrations. The presence of these characteristic bands indicates the successful preparation of Ti-PAN.

**Figure 1.**
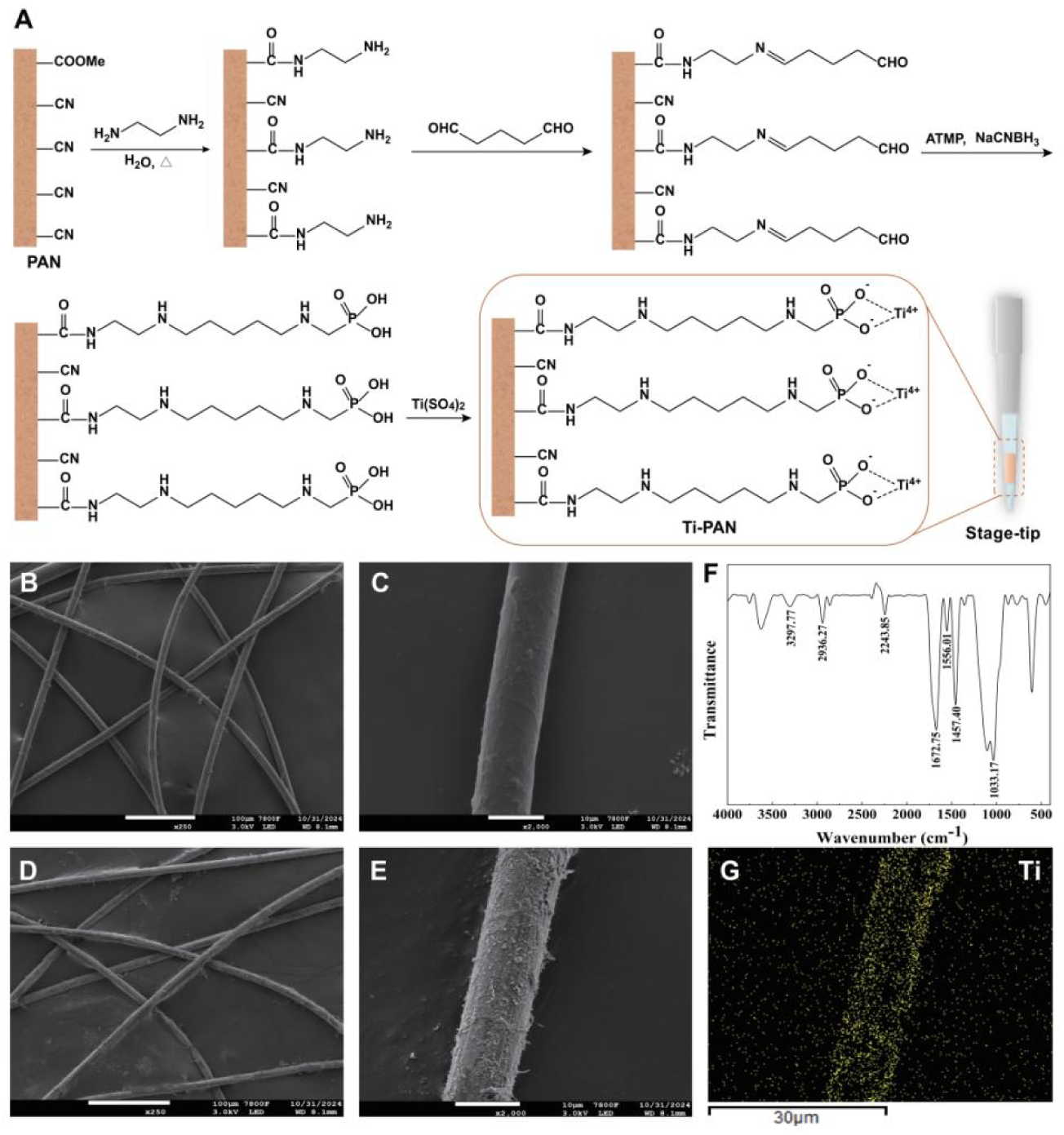
The preparation and characterization of Ti-PAN material (A) Scheme for the preparation of Ti-PAN. The SEM images of PAN (B) ×250, (C) ×2000, Ti-PAN (E) ×250 (D)×2000. (F) The FTIR spectra of Ti-PAN. (G) EDS elemental Ti mapping images of Ti-PAN.

### Evaluating the Phosphopeptide Enrichment Performance of Ti-PAN

The excellent flexibility and permeability of the fibers facilitate easy packing into pipette tips, allowing a straightforward enrichment process via simple aspiration and dispensing. This design not only minimizes sample loss but also enhances analytical sensitivity and is readily amenable to automation for high-throughput applications, which allowing for the simultaneous enrichment of multiple samples using a multichannel pipette or a 96-well plate autosampler. As shown in Figure 2A, the peptide mixture were loaded onto the Ti-PAN tip and washed to remove non-specific bindings, followed by eluting under basic condition to release the retained phosphopeptides. Importantly, each aspirate-dispense cycle takes only 4 seconds, and the entire processing time is completed in just 8 minutes, significantly accelerating phosphopeptide enrichment compared to conventional centrifugation methods^[12, 22, 25–27]^. The phosphopeptide enrichment performance was initially evaluated using a standard phosphopeptide (MAQPFS(phospho)LR) and tryptic digests of a standard phosphoprotein (β-casein) spiked into non-phosphorylated BSA digestion. To enhance enrichment efficiency, the sample loading conditions, specifically the concentrations of lactic acid and ACN in the loading buffer, were optimized (Figures S2 and S3). The results indicated that the highest phosphopeptide capture specificity and the most effective removal of non-phosphorylated peptides were achieved when the loading buffer contained 60% ACN, 6% TFA and 10 mM lactic acid. Remarkably, as shown in Figure S4, the material recovery was evaluated using the standard phosphopeptide, resulting in a high recovery rate of 94.4% (n=3, RSD 12.4%). The enrichment selectivity was further assessed by testing a range of intermediate mass ratios, including standard phosphopeptide-to-BSA digest ratios of 1:10 and 1:100, as well as a β-casein-to-BSA digest ratio of 1:10 (Figure S5). As depicted in Figure 2, when the mass ratio of standard phosphopeptide to BSA digestion was 1:1000, the MALDI-TOF spectrum prior to enrichment was predominantly composed of non-phosphorylated peptides. However, after enrichment with the Ti-PAN tip, most non-phosphorylated peptides were effectively removed, resulting in a significant enhancement of phosphopeptide signals. Notably, the Ti-PAN tip was still able to capture the phosphopeptide effectively even when the interference ratio was increased to 1:5000 (Figure 2). Furthermore, when phosphopeptide enrichment was performed using β-casein digestion, the Ti-PAN tip demonstrated robust interference resistance, maintaining high selectivity with BSA digest ratios up to 1:1000. Taken together, these results underscore the exceptional phosphopeptide enrichment selectivity of the Ti-PAN tip, highlighting its promising application for isolating and enriching phosphopeptides from complex biological matrices.

**Figure 2.**
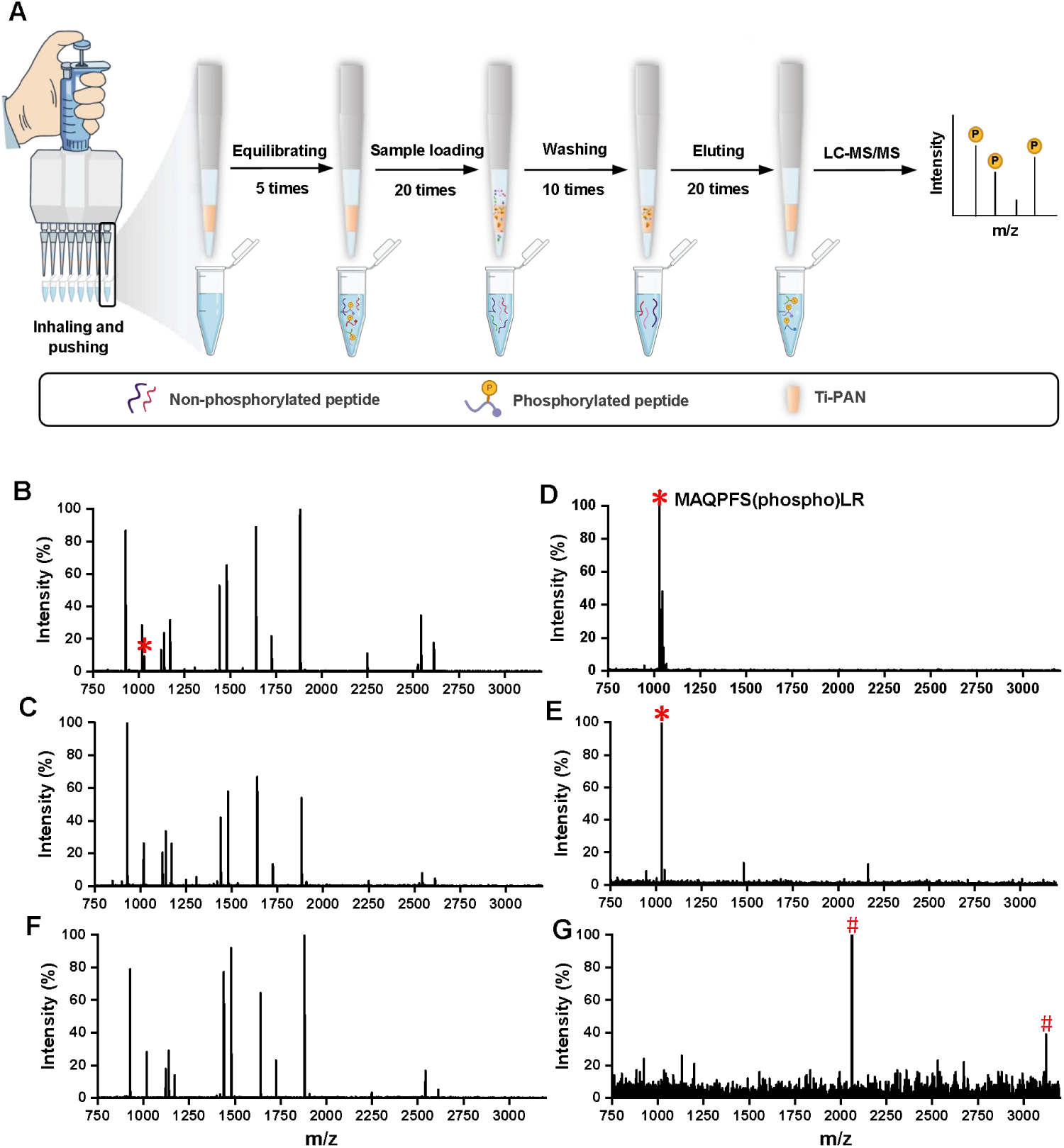
The performance of Ti-PAN on the phosphopeptide enrichment from the mixture of BSA and standard peptide or β-casein. (A) Workflow of phosphopeptide enrichment by Ti-PAN tip. MALDI-TOF MS spectra of the mixture of BSA and (B-E) standard phosphopeptide or (F, G) β-casein at different molar ratios. Direct analysis at (B, F) 1000:1 and (C) 5000:1 ratio. Postenrichment at ratios of (D, G) 1000:1 and (E) 5000:1 ratio. Phosphopeptides are marked with red “*” of tryptic digests from standard peptide and red “#” of β-casein.

### Phosphoproteome profiling of Nanoscale HeLa Cell Samples

Beyond its high selectivity for phosphopeptide enrichment, the Ti-PAN tip allows the entire enrichment process to be completed in-situ within one tip, eliminating transfer steps and thereby reducing loss while boosting analytical sensitivity. We evaluated its performance in phosphoproteomic profiling across low-input cells ranging from 100 to 100,000 cells. As shown in Figure 3A, nanoscale cell suspensions were lysed with dodecyl maltoside (DDM). The lysates were then rapidly digested with trypsin within 2 h, followed by phosphopeptide enrichment using a single Ti-PAN tip in one preparation tube. Finally, the enriched phosphopeptides were analyzed using our recently developed MS acquisition method turboDDA, which has shown exceptional performance for microsample analysis. To enhance sample preparation efficiency, we optimized the concentration of DDM used for lysis of microscale cell samples (1,000 cells). The results indicated that a DDM concentration of 0.01% (wt/v) yielded the highest numbers of identified phosphosites, phosphopeptides, and phosphoproteins (Figure S6).

**Figure 3.**
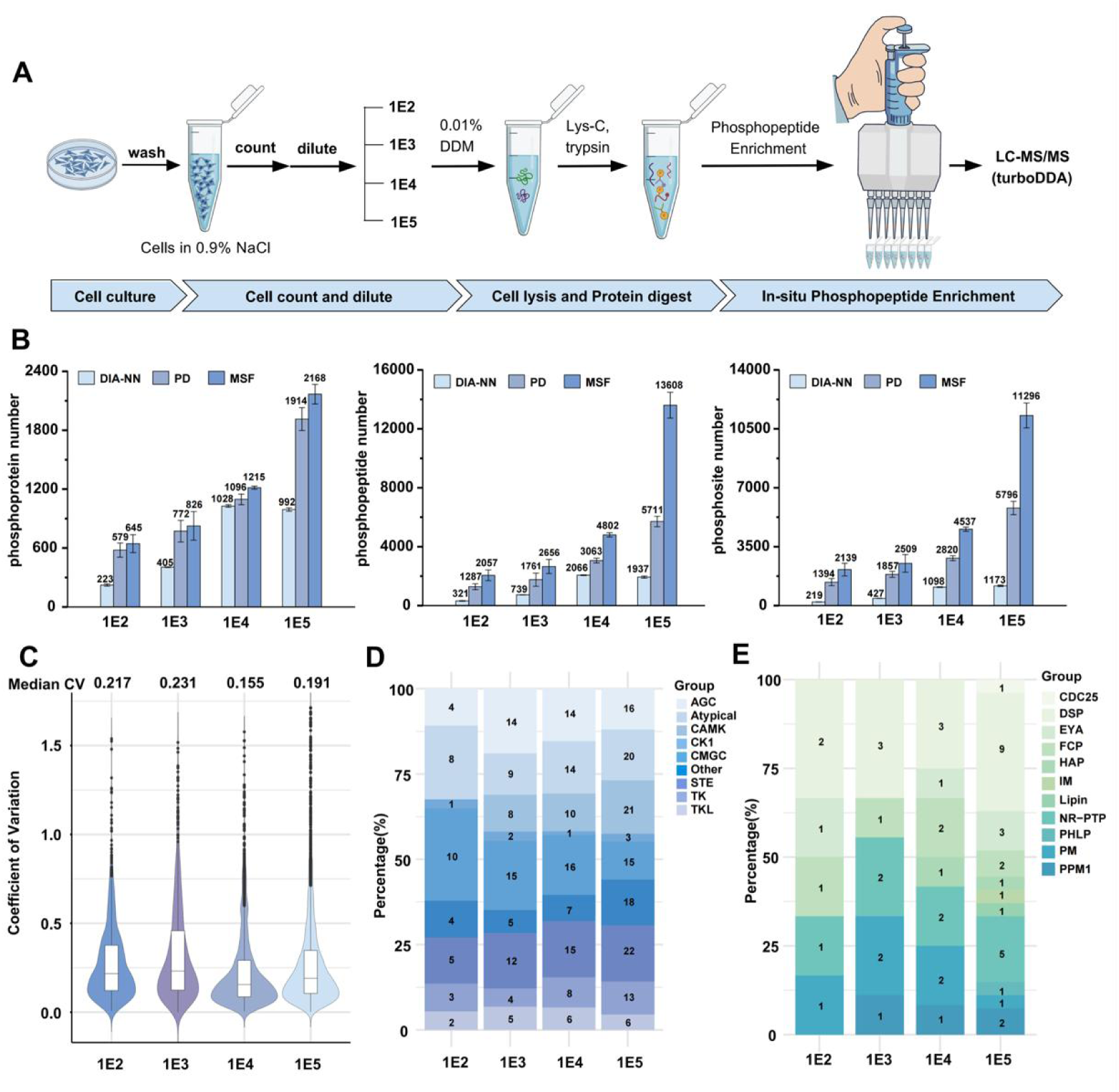
Phosphopeptide enrichment performance of Ti-PAN tip using different amount of HeLa cell as inputs. (A) Phosphopeptide enrichment workflow for the analysis of different amount of HeLa cells. (B) The number of identified phosphoproteins, phosphopeptides, and phosphosites from 1E2 to 1E5 Hela cells via DIA-NN, Proteome discoverer and MSFragger. (C) Coefficient of variation distribution of the LFQ intensity of phosphopeptides identified from 1E2 to 1E5 HeLa cells. (D) Kinase identification number in different groups with different amount of HeLa cells. (E) Phosphatase identification number in different groups with different amount of HeLa cells.

All experiments were performed in triplicate and in parallel, with good consistency and overlap observed among the replicates, as shown in Figure S7. Using Ti-PAN, a substantial number of phosphosites, phosphopeptides, and phosphoproteins were identified from small amounts of HeLa cell digests by both Proteome Discvoerer and MSFragger (Figure 3B). Specifically, from just 100 HeLa cells, 1,394/2,139 phosphosites, 1,287/2,057 phosphopeptides, and 579/645 phosphoproteins were identified with Proteome Discoverer and MSfragger, respectively (Table S1). The number of identifications increased with larger starting cell amounts: from 1,000 cells, 1857/2509 phosphosites, 1761/2656 phosphopeptides, and 772/826 phosphoproteins were detected (Table S2); from 10,000 cells, 2,820/4,537 phosphosites, 3,063/4,802 phosphopeptides, and 1,096/1,215 phosphoproteins (Table S3); and from 100,000 cells, 5,796/11,296 phosphosites, 5,711/13,608 phosphopeptides, and 1,914/2,168 phosphoproteins (Table S4). Notably, MSFragger consistently identified more phosphosites and phosphopeptides than Proteome Discoverer. The fold increase in identifications for 100, 1,000, 10,000, and 100,000 cells was 1.5, 1.35, 1.6, and 1.9, respectively, indicating that the improvements were especially pronounced with higher cell input. Additionally, MSfragger demonstrated clear advantages in both identification numbers and processing time for phosphoproteomic analysis, consistent with previous reports.^[22]^ Furthermore, compared to the DIA MS technique commonly used by most researchers, we found that the turboDDA MS method demonstrated a clear advantage in phosphorylation identification in cell samples of varying quantities (Table S5), as previously verified in our work ^[23]^.

To assess the reproducibility of phosphopeptide enrichment in nanoscale samples, we analyzed the distribution of CVs for each pairwise comparison between biological replicates (Figure 3C). The results showed that the CV values remained consistently low across the nanoscale samples, with an average CV of 0.2, indicating high phosphorylation enrichment reproducibility of our methods.

By mapping 522 human kinases from KinMap^[28]^ and 238 human phosphatases from DEPOD^[29]^, a total of 138 kinases with 5,091 phosphosites in the kinase tree and 38 phosphatases with 1,429 phosphosites were identified from micro-scale cell samples (Fig. 3D, E, Table S6). Among these, 35 kinases and 6 phosphatases were detected from as few as 100 cells, and 33 of kinases and 6 phosphatases were identified in at least two orders of magnitude of low-cell inputs.

Furthermore, to assess the performance of Ti-PAN enrichment of phosphorylated peptides in micro samples, we compared it with the commercial Fe-NTA^[25]^ and TiO_2_ beads^[22]^, which are widely used for phosphopeptide enrichment. To this end, we conducted an experiment in which 9 samples of 10,000 HeLa cells were simultaneously treated in parallel and enriched by Ti-PAN, Fe-NTA and TiO_2_ beads, respectively (n=3). As shown in Figure 4A, the Ti-PAN method identified 4,537 phosphosites, 4,802 phosphopeptides, and 1,215 phosphoproteins from 10,000 HeLa cells. In comparison, Fe-NTA enrichment yielded 3,424 phosphosites, 3,757 phosphopeptides, and 1,137 phosphoproteins (Table S7), while TiO₂ enrichment resulted in only 998 phosphosites, 1,061 phosphopeptides, and 452 phosphoproteins (Table S8). These results highlight the superior enrichment efficiency of Ti-PAN for microscale cell samples, along with a significantly shorter processing time (7.7 minutes), compared to Fe-NTA (100 minutes) and TiO₂ (16 minutes). CV distribution analysis of LFQ intensities across three biological replicates, processed with MSFragger and Proteome Discoverer (Figure 4B), further demonstrated that Ti-PAN consistently yielded lower coefficients of variation, indicating superior reproducibility. Additionally, Ti-PAN covered a phosphoprotein abundance range of more than 6 orders of magnitude, broader than that of Fe-NTA and TiO_2_ (Figure 4C). Although many phosphosites were identified by different approaches, each method still revealed a distinct set of phosphosites (Figure S8). In terms of the phosphosite insight investigation, as shown in Figure 4D, all materials exhibited similar enrichment for phosphoserine, phosphothreonine, and phosphotyrosine. Notably, the proportion of phosphothreonine and phosphotyrosine identified from MSFragger data was higher than that from Proteome Discoverer. For example, in Ti-PAN enrichment, 10.6% and 0.3% of threonine and tyrosine sites were identified using Proteome Discoverer, while 21.6% and 5.7% were identified with MSFragger, respectively. Moreover, the choice of enrichment material markedly influenced the capture of mono-versus multi-phosphorylated peptides. As shown in Figure 4E, Ti-PAN demonstrated a strong enrichment bias toward multiphosphorylated peptides, with 43.4% (MSFragger) and 37.2% (Proteome Discoverer) of the identified peptides carrying multiple phosphorylation sites. This is substantially higher than the proportions observed for Fe-NTA (13.6% using MSFragger, 11.8% using Proteome Discoverer) and TiO₂ (5.3% using MSFragger, 3.8% using Proteome Discoverer), underscoring the enhanced capacity of Ti-PAN for enriching complex phosphorylation patterns.

**Figure 4.**
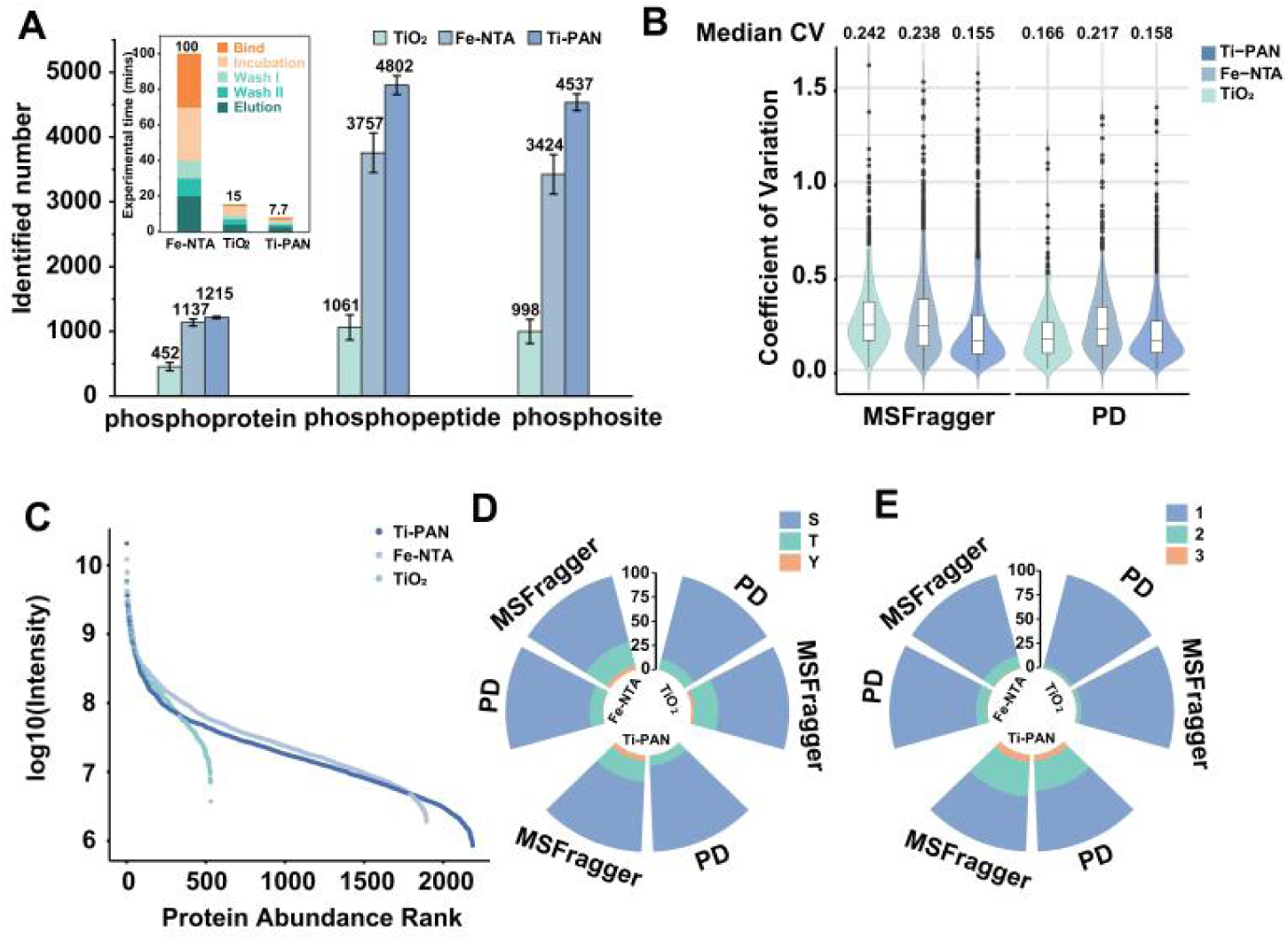
Comparison of the phosphoproteome analysis performance by Ti-PAN, Fe-TNA and TiO_2_. (A) Identified number of phosphoproteins, phosphopeptides and phosphosites by Ti-PAN, Fe-TNA and TiO_2_. The illustration inside is the comparison of phosphopeptide enrichment time for three materials. (B) Coefficient of variation distribution of the LFQ intensity from three replicate of phosphopeptide enrichment experiments by differnt materials. (C) Distribution of the log_10_ intensity for the phosphopeptides identified by three materials. (D) The Serine (S), Threonine (T) and Tyrosine (Y) phosphosites identification number with Proteome Discoverer and MSFragger by three materials (E) The identification number of phosphopeptides with 1, 2, or 3 phosphorylation sites in HeLa cells for three materials.

### Phosphoproteome Analysis of Nanoscale Samples from AP-MS and Mouse Brain Slice

Building on the exceptional performance of Ti-PAN, we further explored its phosphorylation enrichment capabilities in other classical nanosample types, including AP-MS, mouse brain slices, and oocyte samples. AP-MS, a key technique for protein-protein interaction analysis, has traditionally required cell quantities on the order of 10⁸ for in-depth studies, particularly when investigating its phosphorylation. The study of ERK protein interactions is critical for understanding intracellular signal transduction, especially in cell proliferation, differentiation, and survival. Phosphorylation plays a central role in modulating these interactions, as it dynamically regulates protein activity and interaction specificity within the ERK/MAPK signaling pathway. To investigate the ERK interactome, we employed an AP-MS approach combined with label-free quantitative analysis to monitor changes in the MAPK3 interaction network.^[30]^ Following immunoprecipitation, phosphopeptide enrichment was performed to enable in-depth phosphoproteomic profiling of the ERK signaling complex from as few as 10⁵ cells (Figure 5A). The high intra-group correlation coefficients for both the CTRL and ERK groups (>0.9) and low inter-group correlation coefficients (∼0.5), indicated the good quantification reproducibility and high efficient ERK interatome identification (Figure 5B, Table S9). Additionally, the phosphoproteome analysis revealed that 427/499 phosphoprotein, 814/1,461 phosphopeptide and 723/1,396 phosphosite via Proteome Discoverer and MSFragger (Figure 5C), with totally 2,234 phosphosites on 600 phosphoproteins identified. As shown in Figure 5D, a total of 1,113 MAPK3-interacting proteins (upregulated) were identified, including MAPK3 itself and several ERK-interacting proteins (such as MAP2K2, MKNK1, MKNK2, NF1, DUSP9, and RPS6KA3), all of which were significantly upregulated. Among them, 155 proteins were phosphorylated with 632 phosphosites. KEGG analysis of interacting phosphoproteins (Figure 6E) revealed significant enrichment in pathways closely related to key cellular functions, including mTOR, TGF-β, AMPK, MAPK, ErbB, Toll-like receptor, TNF, and EGFR tyrosine kinase inhibitor resistance signaling pathway. Notably, nine ERK-interacting proteins were identified within the MAPK signaling pathway, with seven of these proteins, such as MAPK3, RPS6KA3, MAP2K2, NF1, RRAS2, DUSP9, and RAF1, being phosphorylated at 18 distinct sites (Figure 5F). Additionally, six phosphorylation sites (T198, T202, T207, S110, S283, and Y204) were identified on ERK1, including two new phosphorylation sites S110 and S283. Protein-protein interaction (PPI) network analysis of kinase-substrate interactions from the STRING database revealed that several upregulated kinases, including MAPK3, serve as key nodes interacting with numerous substrates (Figure 5G). Notably, MAPK3 was identified as a central hub, with 30 substrates and 48 phosphorylation sites, indicating its central regulatory role under the experimental conditions. Overall, the use of our highly sensitive phosphorylation enrichment material, Ti-PAN, demonstrates significant potential for functional phosphoproteomic analysis in trace amounts of AP-MS samples.

**Figure 5.**
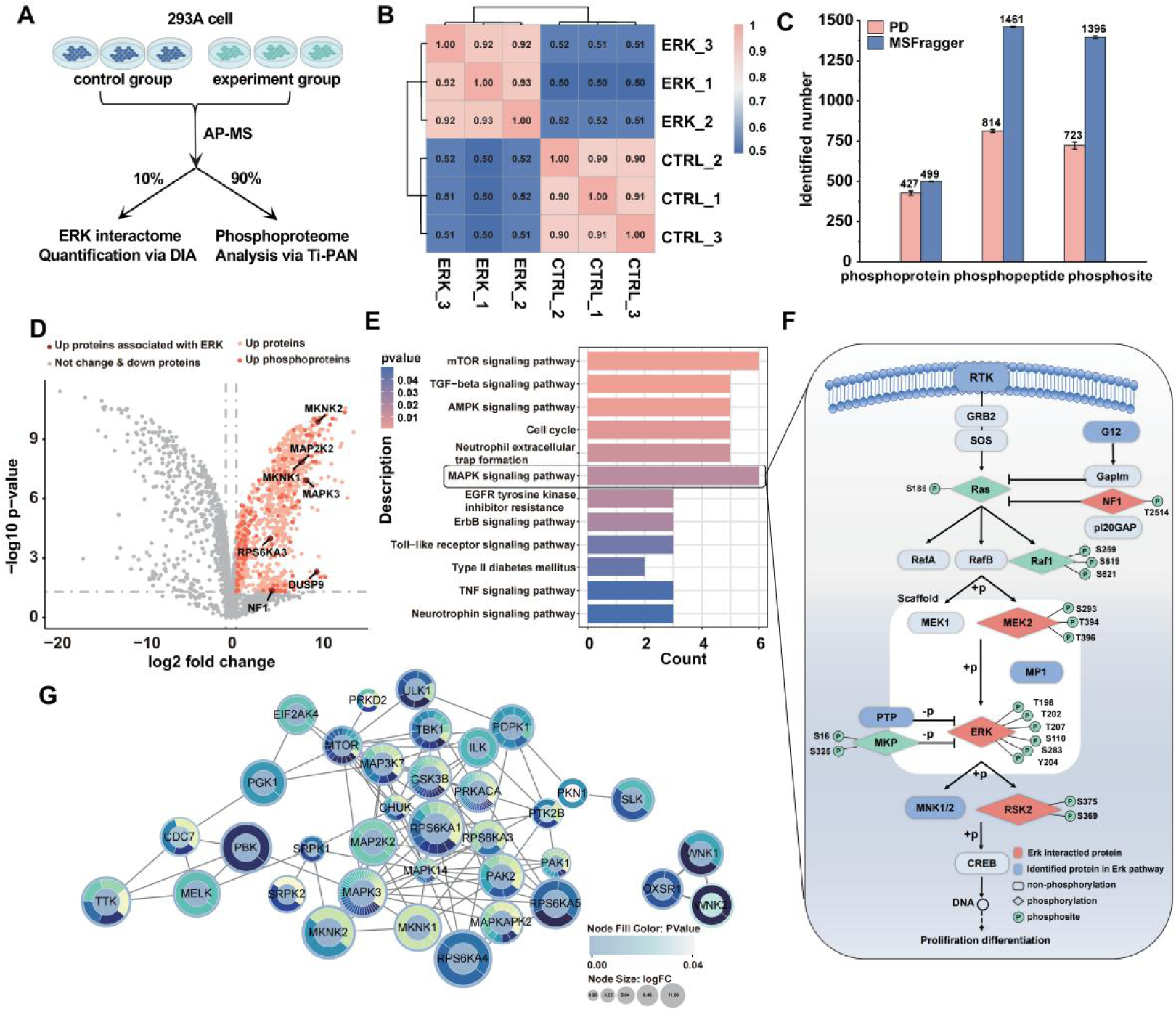
The performance of Ti-PAN tips on the phosphoproteome analysis of AP samples. (A) The workflow to decipher ERK interactome and its phosphorylation by comparing 293A cell lines stably expressing pLVX-MAPK3-GFP-biotin (experiment group) with pLVX-GFP-biotin (control group) proteins. (B) Correlation amon<u>g</u> different replicates. (C) The number of identified phosphoproteins, phosphopeptides, and phosphosites. (D) Volcano plot showing the differentially expressed proteins, phosphoproteins and ERK associated proteins (E) Gene ontology analysis of the biological processes of differential expression proteins. (F) Pathway map of MAPK3 signaling-related proteins. (G) Kinase-substrate interaction network analysis, the node size indicates the fold change in expression and the color intensity represents the number of substrates.

**Figure 6.**
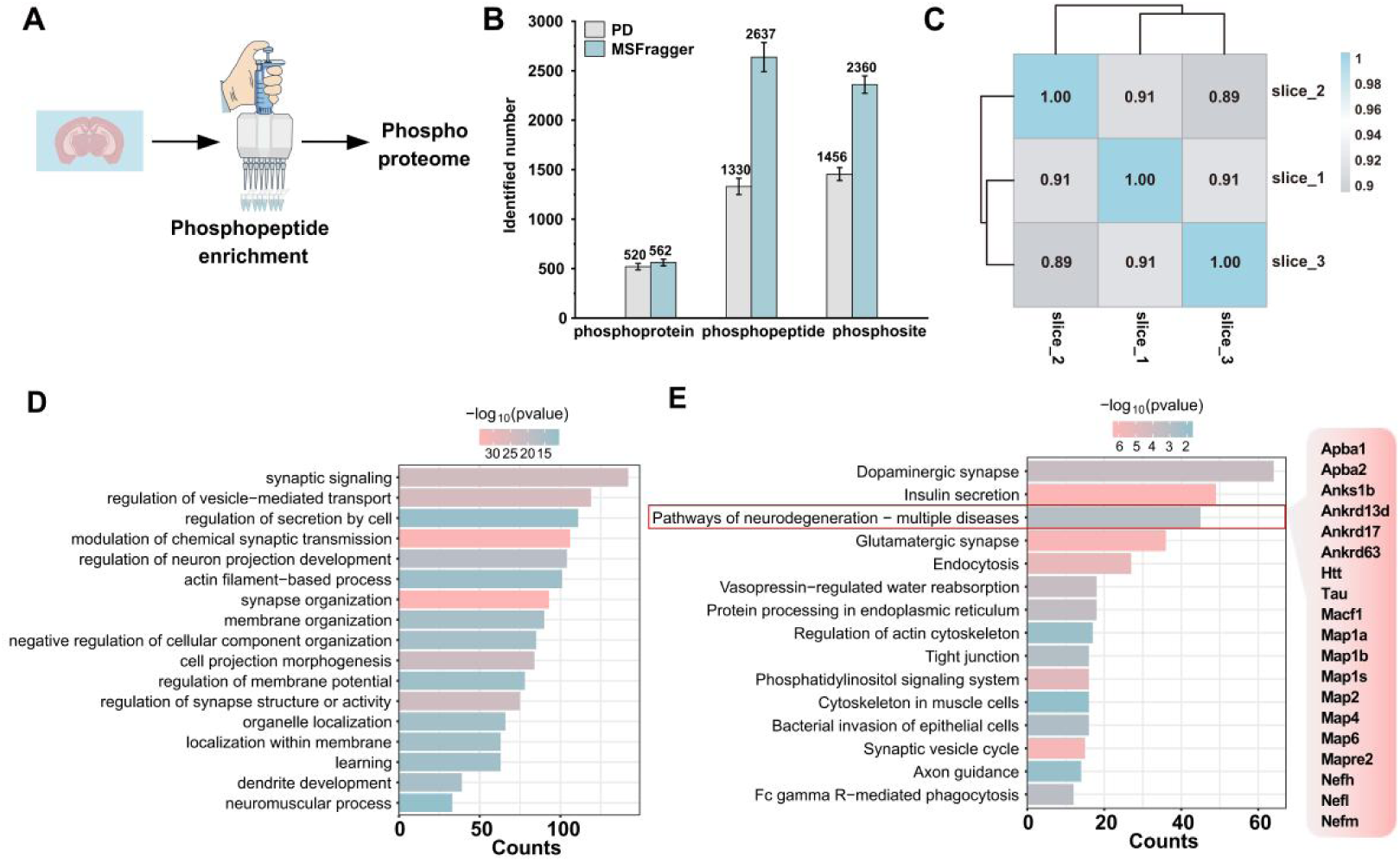
The performance of Ti-PAN tips on the phosphoproteome analysis of low input mouse brain slice (A) The procedure of phosphopeptide enrichment of mouse brain slice by Ti-PAN tips. (B) The number of identified phosphoproteins, phosphopeptides, and phosphosites from low input mouse brain slices via Proteome Discoverer and MSFragger. (C) Correlation among different replicates. (D) Bar graph showing the statistically enriched GO pathways in the phosphoproteome. (E) Bar graph showing the statistically enriched KEGG pathways in the phosphoproteome. Benjamini-Hochberg (BH) corrected P value adjustment (P.adjust) was used for the enrichment analyses.

The analysis of the phosphoproteome in mouse brain tissue slices plays a crucial role in advancing spatial proteomics by integrating cellular signaling into the study of tissue architecture. By extending phosphoproteomic analysis to these tissue slices, we can explore how signaling networks function within their native spatial and cellular contexts. However, achieving in-depth phosphoproteomic analysis from the limited material of such tissue sections remains a significant challenge, underscoring the need for the development of highly sensitive phosphoproteomics technologies. In the study, a total of 2,360±79 phosphosites, 2,637±131 phosphopeptides, and 562±33 phosphoproteins identified from 10 μm mouse brain slices (n=3), contributing valuable insights into the phosphorylation patterns that regulate various neural processes (Figure 6A and 6B, Table S10). The reproducibility of the analysis was confirmed by the high correlation among different technical replicates, as shown in Figure 6C, which indicated a high degree of consistency across multiple experimental runs. The study also examined biological processes (BP) and Kyoto Encyclopedia of Genes and Genomes (KEGG) pathways related to the identified phosphoproteins (Figure 6D and 6E). Notably, the phosphoproteins were enriched in pathways related to synaptic transmission, neurogenesis, and cellular signaling, which are critical for maintaining normal brain function and neuroplasticity. Furthermore, several proteins associated with neurodegenerative diseases, such as tau and amyloid-beta, were identified within the dataset. These proteins are implicated in conditions like Alzheimer’s and Parkinson’s disease, where abnormal phosphorylation and aggregation play key roles in disease progression. Some AD marker proteins, such as Tau (P10637), 32 phophosites were identified, among which 29 sites (including two phosphotyrosine sites Y489 and Y686) were consistent with the reported and 3 sites (Thr 60, Thr 65 and Thr 678) were newly identified. 5, 19 and 49 phosphosites were identified from NFL (P08551), NFM (P08553) and NFH (P19246), respectively. Among the NFH protein, 47 sites have been reported, 2 sites (S63 and S597) were newly identified. By identifying and mapping phosphorylation sites on proteins critically involved in neurodegenerative diseases, such as Tau and NFH, this proteomic dataset pinpoints specific molecular mechanisms and newly discovered regulatory sites that could serve as potential targets for therapeutic intervention.

### Phosphorylation Dynamics Analysis During Mouse Oocyte Maturation

Oocytes play a critical role in female reproduction, and identifying oocyte-specific phosphorylation sites is essential for understanding their specialized developmental program. However, effective analysis of the phosphorylation dynamics in mammalian oocyte remains challenging, owing to the scarcity of oocyte samples and the absence of high-efficiency enrichment methods. In this study, as illustrated in Figure 7A, we applied a high-sensitivity phosphopeptide enrichment strategy combined with quantitative MS-based phosphoproteomics to characterize the phosphorylation dynamics of mouse oocytes during maturation. This was achieved by isolating just five oocytes at four key time points: GV (germinal vesicle) stage, meiotic resumption at GVBD (germinal vesicle breakdown), MI (metaphase I) , and ovulated MII (metaphase II) stage (with 5 oocytes per stage and 5 replicates). As shown in Figure 7B (Table S11), 174±31, 417±20, 534±92, and 1068±235 phosphosites (corresponding to 88±12, 184±5, 241±73, and 402±67 phosphoproteins, as well as 190±28, 448±22, 568±102, and 1197±214 phosphopeptides) were identified from oocytes at the GV, GVBD, MI, and MII stages, respectively. In total, we achieved high-quality quantification of 2709 distinct phosphosites (with >0.75 localization probability) located on 701 proteins from mouse oocytes, indicating that an average of 27 phosphosites per cell could be obtained, which represents the deepest coverage of phosphoproteomes from such minimal oocyte material reported to date. Traditionally, tens thousands of oocytes were needed for the phopshoproteome analysis^[31]^. Additionally, 536 phosphosites (19.8%) and 115 phosphoproteins (16.4%) were commonly identified across all four stages (Figure 7C). Notably, when compared with the PhosphoSitePlus database^[32]^, we found that 44.9% (1,217 out of 2,709) of phosphosites on 423 proteins in oocytes were undetected in other cell types (Figure 7D). Correlation analysis results (Figure 7E) show that biological replicates of mouse oocytes within the same stage exhibit a high degree of correlation, with their respective coefficients of variation (CV) being less than 5% (Figure S9), demonstrating the high reproducibility of the analysis method. Additionally, the correlation of phosphorylation in mouse oocytes across different stage is relatively low, and principal component analysis (PCA) analysis reveals clear clustering of oocytes from the four stages, indicating their distinct differences (Figure 7F). Therefore, analyzing the phosphorylation heterogeneity across various stages is crucial for understanding the dynamic regulatory mechanisms that govern the maturation process of mouse oocytes. Quantitative phosphoproteomic analysis revealed that, compared with GV stage oocytes, the GVBD, MI, and MII stages exhibited 187, 263, and 487 upregulated phosphosites and 28, 28, and 39 downregulated phosphosites, respectively (Figure S10). Furthermore, clustering analysis was performed on the 678 dysregulated phosphosites (Figure 7G), which were categorized into four major clusters based on their dynamic phosphorylation patterns during oocyte maturation. Cluster 1 showed a transient upregulation at the GVBD stage, followed by a decrease, and was primarily enriched in processes related to mRNA stabilization and translation regulation. Cluster 2 displayed a progressive increase from GV to MII, with enrichment in chromosome segregation and spindle organization. Cluster 3 exhibited increased phosphorylation during MI, and was associated with organelle assembly and actin cytoskeleton organization. In contrast, Cluster 4 showed a sharp increase at the MII stage, enriched in pathways involved in cell cycle progression, spindle organization, and translational initiation. These findings suggest that distinct phosphosite clusters coordinate different biological processes during oocyte meiotic maturation.

**Figure 7.**
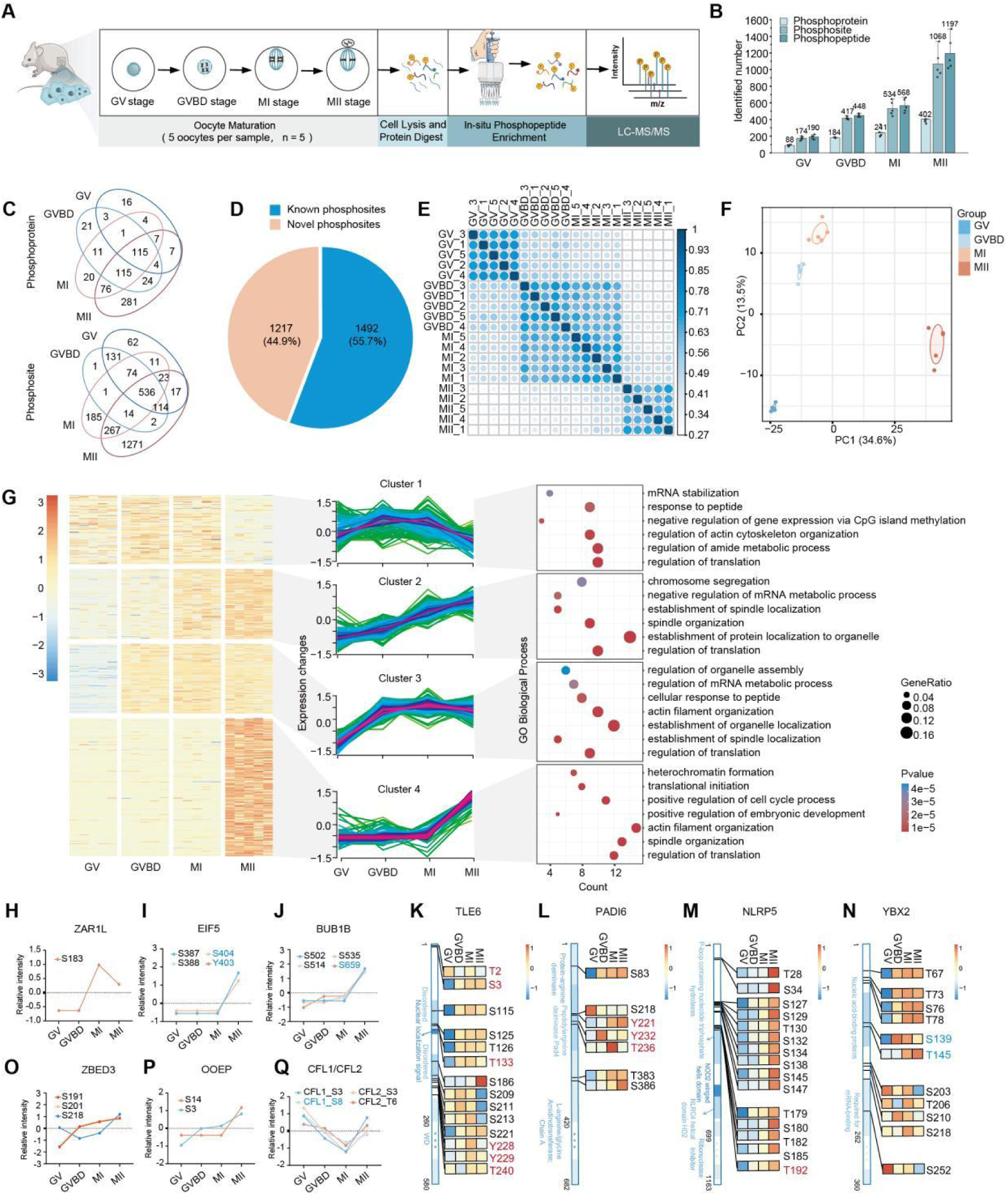
Phosphorylation dynamic of mouse oocytes during meiotic maturation. (A) Workflow of the phosphopreoteome analysis of mouse oocytes at GV, GVBD, MI and MII stage by Ti-PAN. (B) Number of identified phosphoproteins, phosphopeptides and phosphosites at different stages. (C) Overlap of the identifed phosphoproteins and phosphosites from different stages. (D) The pie chart showing the novel and known phosphosites by comparing the results in our study and PhosphoSitePlus database. (E) Correlation among the samples from different stages. (F) Principal component analysis depicting the clustering of five phosphoproteome replicates from GV, GVBD, MI and MII oocytes. (G) Heatmap illustrating the dynamic changes in regulated phosphosites during oocyte maturation stages. Fuzzy c-means clustering organized the phosphosites into four distinct clusters. Each line indicates the relative abundance of individual phosphosites. Representative biological processes for each cluster are show on the right. (H-P) Relative abundance of phosphosites in representative proteins at different stages, with red and blue colors denoting sites newly identified in this study and those not previously detected in mouse oocytes, respectively. (K-N) Location of the identified phosphosites within their respective protein domains. Red and blue colors denoting sites newly identified in this study and those not previously detected in mouse oocytes, respectively.

Further zooming in on each phosphosite, we observed that many sites exhibited phosphorylation level trends during mouse oocyte maturation that are consistent with previously reported findings. For example, as shown in Figure S11, phosphorylation at MAPK3 (Q63844, T203), MAPK1 (P63085, T183 and Y185), andDNMT1 (P13864, S1128 and S331), suggesting that our method provides accurate quantification. Interestingly, we also found some new phosphosites exhibited significant changes. For instance, ZAR1L and its homolog ZAR1 regulatedthe maternal transcriptome and translational activation in maturing oocytes.^[33]^ We found the ZAR1L phosphorylation at S183 was significantly upregulation in MI stage (Figure 7H). EIF5 (P59325), a key translation initiation factor in oocytes, is essential for oocyte growth and meiotic maturation^[34]^. The translational activity of EIF5 depends on its phosphorylation status, and casein kinase II (CKII) acts as its upstream kinase^[35]^. Consistent with this, inhibition of CKII activity in Xenopus oocytes also leads to spindle formation defects^[36]^. In our data, phosphorylation of EIF5 at residues S387, S388, S404, and Y403 displayed stage-specific dynamics during oocyte maturation. While phosphorylation levels at these sites were low at the GV and GVBD stages but increased significantly upon progression to MI and MII (Figure 7I).

This phosphorylation pattern correlates temporally with the period of spindle organization, indicating its role in activating EIF5 for translation initiation. Another interesting protein is BUBR1 (Q9Z1S0), a spindle assembly checkpoint protein required for faithful chromosome segregation during meiosis^[37]^. Studies in Xenopus oocytes demonstrated that loss of BubR1 impairs the interaction between Mad2, Bub3, and Cdc20, an anaphase activator. Although BubR1 phosphorylation is not strictly required for recruiting all other checkpoint components to kinetochores, its own stable kinetochore localization and characteristic hyperphosphorylation were found to be dependent on Bub1 and Mad1^[38]^. Phosphorylation at S670 has been shown to be essential for error correction and for kinetochores with end-on microtubule attachments to establish tension and the phosphorylation status of BubR1 is important for checkpoint-mediated inhibition of the anaphase-promoting complex/cyclosome (APC/C)^[39]^. Our data reveal that BUBR1 phosphorylation at S502, S514, S535, and S659 are dynamically regulated during mouse oocyte maturation (Figure 7J). This pattern aligns with the role of BUB1R in regulating the spindle assembly checkpoint and meiotic progression, suggesting that these modifications may contribute to its full functional engagement during meiotic stages (Figure 7J).

Notably, many of the altered phosphorylation sites we detected in oocytes mapped to components of the subcortical maternal complex (SCMC) complex. The SCMC exhibits a distinctive localization in the subcortical regions of oocytes and pre-implantation embryos and is associated with crucial pathways during the transition of an oocyte to an embryo.^[40]^ Our data revealed several new phosphosites within their functional domains (Figure 7K-Q). TLE6 deficient oocyte exhibits asymmetric cleavage of the post-fertilization embryo, resulting in the developmental arrest.^[41]^ Our profiling identified 14 distinct phosphosites changed during oocyte maturation, 7 of which have not been previously reported. Similarly, PADI6 deficiency disrupts the formation of cytoplasmic lattices in mouse oocytes^[42]^, while in human, loss-of-function maternal-effect variants of PADI6 are associated with multi-locus imprinting disturbances with heterogeneous manifestations in offspring.^[43]^ We also discovered 3 novel phosphosites (Y221, Y232 and Y236) within the core domain of PADI6, whose phosphorylation levels increased following GVBD (Figure7L). Additionally, other substantial novel phosphorylation sites were identified in SCMC component proteins, including NLRP5-T130, T192, YBX2-S139, T145, S252, ZBED3-S191, S218, and OOEP-S14 (Figure7M-7Q). These widespread phosphorylation changes warrant further investigation as potential regulatory mechanisms regulating the quality of oocyte and early embryonic development. Moreover, our phosphorylation landscape may provide a molecular resource for elucidating the etiology of maternal-effect genetic disorders.

### Validation of Kinase Involvement in Oocyte Maturation

Phosphorylation of key proteins is essential for oocyte maturation. We performed kinase prediction based on upregulated phosphorylation sites at various stages (GVBD, MⅠ and MⅡ) relative to the GV stage (Figure S10), thereby uncovering key kinases potentially involved in regulating oocyte maturation. Through the kinase analyses, 52 kinases (termed predicted kinases here) was estimated according to the phosphorylation changes in their known substrates and 19 kinase was identified in our MS data (termed identified kinases here) (Figure 8A and 8B). These kinases belong to all major kinase families, with a higher representation of CMGC family (Fig 8A and 8B). Notably, TTK and MAPK3 were among the predicted kinases that were also identified (Figure 8C and 8D). It has been proven that TTK/MPS1 is a key kinase that safeguards centromeric cohesin protection by recruiting Sgo2^[44]^ and that promotes timely spindle bipolarization before kinetochores stably attach to microtubules^[45]^. MAPK3/ERK1 was an essential kinase for the maintenance of the MII spindle. ERK1/2-deleted oocytes had poorly-assembled MII spindles, spontaneously released polar body-2 (PB2), and were arrested at another metaphase called metaphase III (MIII)^[46]^. We listed the kinases predicted from the phosphorylation sites upregulated in each stage relative to the GV stage in the heatmap (Figure 8E). Among them, many kinases have been demonstrated to play roles in meiotic resumption and oocyte maturation in mice, such as GSK3A^[47]^ from the CMGC family, WEE2^[11, 48]^, and Zap70^[49]^ from the TK family (Figure 8B). In particular, we examined the impact of STK4 (serine/threonine kinase 4) and DYRK3 (dual specificity tyrosine phosphorylation regulated kinase 3) inhibition on oocyte maturation. The fully grown GV oocytes were cultured in medium supplemented with various concentrations of STK4 inhibitor (SBP-3264), and maturation-related phenotypes were evaluated at the 4 h and 16 h after release (Figure 8F). The results showed that SBP-3264 treatment prevented oocytes from undergoing normal GVBD within 4 h, especially at high concentrations (50 μM). Even after extending the culture period to 16 h, GVBD remained suppressed. At a lower concentration (20 μM), approximately 45% of oocytes underwent GVBD by 16 h, while 27.5% degenerated (Figure 8G and 8H). Immunofluorescence analysis further revealed that SBP-3264-treated oocytes failed to form the MI spindle even after 16 h of culture (Figure 8I). The DYRK3 inhibitor GSK626616 was also co-cultured with fully grown oocytes to assess its impact on meiotic progression. Treatment with GSK626616 resulted in decreased rates of GVBD and Pb1 extrusion at 4 and 16 hrs after release (Figure 8K). At higher inhibitor concentrations (50 μM), all oocytes remained arrested at the GV stage even after 16 h of culture, while lower concentrations allowed partial GVBD and Pb1 extrusion but with increased rates of degeneration (Figure 8L). Immunofluorescence analysis further demonstrated that oocytes treated with GSK626616 exhibited disrupted MII spindle assembly and aberrant chromosome alignment (Figure 8M and 8N). These results indicate that STK4 plays an essential role in meiotic resumption in oocytes and DYRK3 is functionally required for timely meiotic resumption and normal maturation in mouse oocytes.

**Figure 8.**
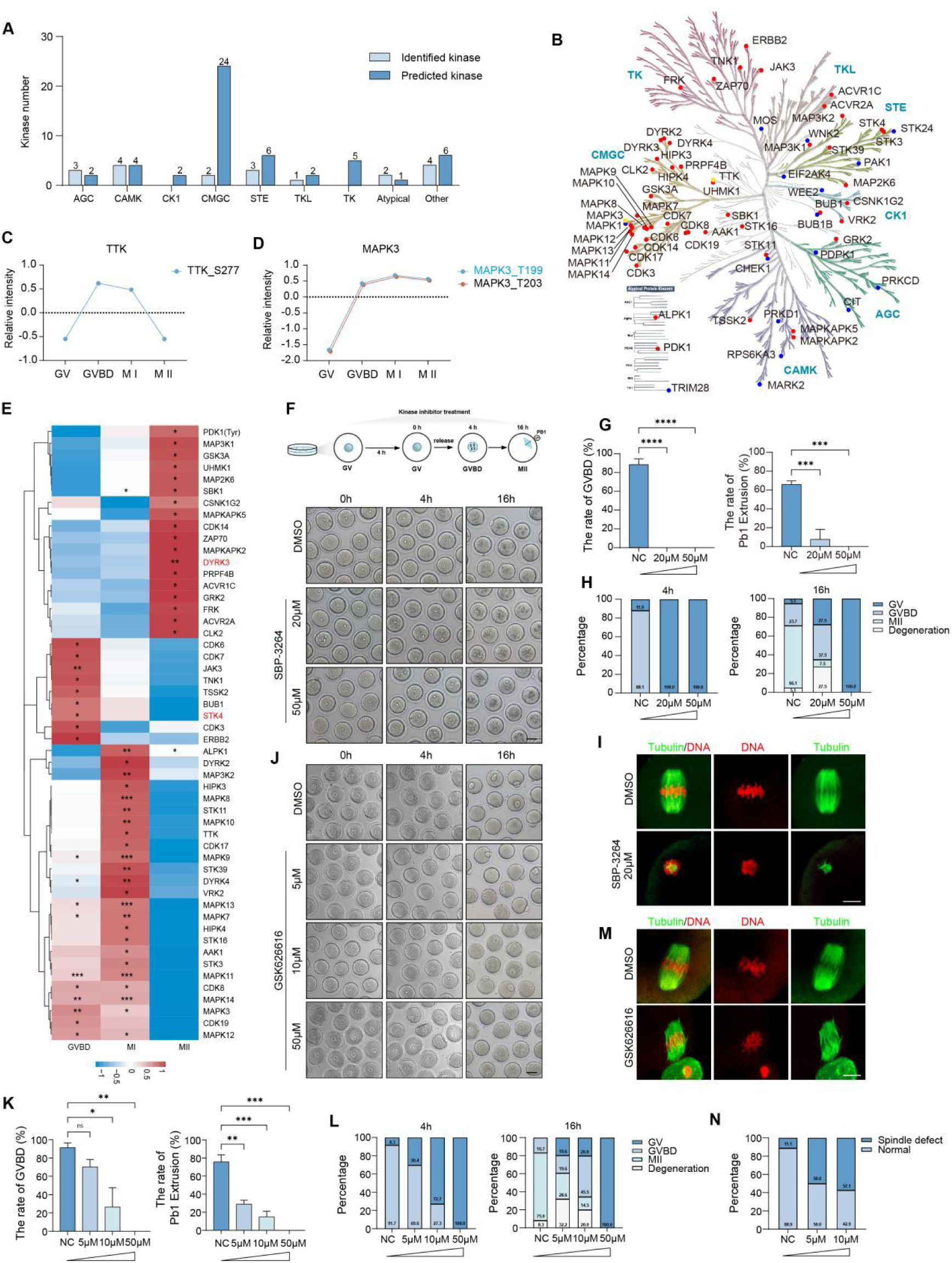
Identification of critical phosphorylated kinases in mouse oocyte maturation. (A) Bar plot representing the number of phosphorylated kinases that identified regulated or predicted in oocytes from each of the major mammalian kinase families. (B) Kinase tree showing the identified (blue) and predicted (red) kinases annotated to the major kinase families. Yellow dote means both (C-D) The change in 4 indicated oocyte stage identified regulated phosphosites on TTK (C), MAPK3(D), with blue colors denoting sites not previously detected in mouse oocytes, (E) Heatmap displays the predicted phosphorylation status of protein kinases at each indicated stage, in comparison to the GV stage. Phosphorylation sites showing significant upregulation in three comparative stages were analyzed against non-altered phosphosites via Fisher’s exact test. The asterisks denote statistical significance: ***p < 0.001, **p < 0.01, *p < 0.05. (F) Bright-field images of control and SBP-3264 treated oocytes at 0 h, 4 h and 16 h after release. Scale bars, 50 μm. (G) The lift bar plot representing the rate of GVBD in control and SBP-3264 treated oocytes at 4 h after release. The right bar plot representing the rate of PB1 extrusion in control and SBP-3264 treated oocytes at 16 h after release. Data are expressed as mean percentage ± SD (n=133). Two-tailed Student’s t test was used for statistical analysis. (H) Bar plot showing the proportion of control and SBP-3264-treated oocytes at various stages (GV, GVBD and MⅡ) or in a degenerated state at 4 h (lift panel) and 16 h (right panel) after release, respectively (n=133). (I) Immunofluorescence images show α-tubulin (green) and chromosomes (red) in control and SBP-3264-treated oocytes at 16 h after release. Scale bars, 10 μm. (J) Bright-field images of control and GSK626616-treated oocytes at 0 h, 4 h and 16 h after release. Scale bars, 50 μm. (K) The lift bar plot representing the rate of GVBD in control and GSK626616 treated oocytes at 4 h after release. The right bar plot representing the rate of PB1 extrusion in control and GSK626616 treated oocytes at 16 h after release. Data are expressed as mean percentage ± SD (n=180). Two-tailed Student’s t test was used for statistical analysis. (L) Bar plot showing the proportion of control and GSK626616-treated oocytes at various stages (GV, GVBD and MⅡ) or in a degenerated state at 4 h (lift panel) and 16 h (right panel) after release, respectively (n=180). (M) Immunofluorescence images show α-tubulin (green) and chromosomes (red) in control and GSK626616 -treated oocytes at 16 h after release. Scale bars, 10 μm. (L) Bar plot showing the proportion of control and GSK626616-treated oocytes at various stages (GV, GVBD and MⅡ) or in a degenerated state at 4 h (lift panel) and 16 h (right panel) after release, respectively (n=180). (N) Bar plot showing the percentage of oocytes with spindle defects in control and GSK626616-treated groups.

All the above results reveal that this approach provides valuable insights into how phosphorylation events influence key processes like oocyte maturation, meiotic progression, and the acquisition of developmental competence. It also helps identify critical phosphorylation sites that could serve as biomarkers or therapeutic targets for improving fertility treatments and understanding fertility-related disorders.

### Conclusions

In conclusion, we have developed a novel Ti-PAN-based phosphopeptide enrichment method that significantly enhances the sensitivity and efficiency of phosphoproteome analysis, especially for microscale samples. By utilizing highly hydrophilic textile PAN fibers in a simple, rapid, and automation-friendly design, our approach allows for the efficient in-situ enrichment of phosphopeptides, minimizing sample loss and improving analytical accuracy. The integration of a rapid tryptic digestion workflow further enhances the method’s applicability to a variety of biological samples. Our results demonstrate the robustness of the Ti-PAN tip, which enabled the identification of thousands of phosphosites across a wide range of sample sizes, from single cells to mouse oocytes, providing unprecedented coverage of the phosphoproteome. Notably, several key kinases involved in meiotic resumption and oocyte maturation were identified, including STK4 and DYRK3. This method has great potential for advancing phosphoproteomic research, offering deep insights into cellular signaling and developmental processes with minimal sample input. Its versatility and high throughput capability make it an invaluable tool for exploring complex biological systems, including those with limited sample availability, such as oocytes and embryos, and could lead to further discoveries in developmental biology and disease mechanisms.

## Data availability

The mass spectrometry proteomics data have been deposited to the ProteomeXchange Consortium (https://proteomecentral.proteomexchange.org) via the iProX partner repository with the dataset identifier PXD071174.

## Supporting Information

Supporting Information is available from …….

## Acknowledgements

This project was supported by the Scientific Research Foundation of the Higher Education Institutions of Liaoning Province China (No. LJ212410165049), the Natural Science Foundation of Liaoning Province, China (No. 2025-MSLH-438), the Liaoning Normal University Doctoral Research Start-up Fund (No. 2024BSL002), the National Natural Science Foundation of China (NO.22574175; 82471693), a Major Scientific Program of CITIC Group (No. 2023ZXKYB34100), the Hunan Provincial Grant for Innovative Province Construction (2019SK4012), the Natural Science Foundation of Hunan Province (No. 2024JJ4100), the Scientific Research Foundation of Reproductive and Genetic Hospital of CITIC-XIANGYA (YNXM-202313; YNXM-202319) and a Hundred Youth Talents Program of Hunan Province.

