## Supplemental file for "Unlocking Ultrasensitive Phosphoproteomics of Nanoscale Samples: A Rapid IMAC Enrichment Platform Using Ti-PAN Tip"

### Table of Contents


| Description | Page No. |
|---|---|
| **Supplementary Experiments** | 3 |
| **Figure S1.** EDX spectrum and chemical composition (table) of Ti-PAN. | 10 |
| **Figure S2.** MALDI-TOF mass spectra of β-casein and BSA tryptic digest mixtures before and after enrichment by Ti-PAN with different loading buffer solutions with various lactic acid concentration | 11 |
| **Figure S3.** MALDI-TOF mass spectra of β-casein and BSA tryptic digest mixtures after enrichment by Ti-PAN with different loading buffer solutions with various ACN concentration | 12 |
| **Figure S4.** Evaluation of the phosphopeptide (MAQPFpSLR) recovery on the PAN-tip. | 13 |
| **Figure S5.** MALDI-TOF MS spectra of the mixture of BSA and standard peptide or β-casein at different molar ratios. | 14 |
| **Figure S6.** The number of identified phosphoproteins, phosphopeptides, and phosphosites at different concentrations of DDM. | 15 |
| **Figure S7.** The Venn plots of identified phosphoproteins and phosphopeptides number from 1E2 to 1E5 Hela cells via Proteome Discoverer and MSFragger. | 16 |
| **Figure S8.** The Venn plots of identified phosphoproteins, phosphosites and phosphopeptides from 10,000 Hela cells with different materials via MSFragger. | 17 |
| **Figure S9.** Coefficient of variation distribution of the LFQ intensity from five replicates at different stages. | 18 |
| **Figure S10.** Volcano plots showing the differential phosphorylated proteins between GVBD and GV, or between MI and GV, or between MII and GV. | 19 |

| | |
|---|---|
| **Figure S11.** Relative abundance of phosphosites in representative proteins at different stages. | 20 |
| **Table S1-5**. List of identified phosphoproteins, phosphosites and phosphopeptides from 1E2 to 1E5 HeLa cells. | |
| **Table S6**. List of identified phosphosites of phosphatases and kinase from 1E2 to 1E5 HeLa cells | |
| **Table S7-8**. List of identified phosphoproteome from 1E4 HeLa cells via $TiO_2$ and Fe-NTA materials | |
| **Table S9.** List of identified phosphoproteome of ERK protein interatome from the AP-MS sample. | |
| **Table S10**. List of quantified phosphoproteome from the mouse brain slice sample. | |
| **Table S11.** List of phosphoproteome from the mouse oocytes. | |

### Supplementary Experiments

**Chemicals and Materials.** The Polyacrylonitrile fiber (PAN, average Mw 150,000) was obtained from Macklin. Ethylenediamine, glutaric dialdehyde (50% in $H_2O$), triethylammonium bicarbonate (TEAB, pH 8.5), acetonitrile (ACN), lactic acid (~85%), (aminomethyl) phosphonic acid (AMPA, 99%), sodium cyanoborohydride ($NaCNBH_3$), ammonia (28~30%), iodoacetamide (IAA), dithiothreitol (DTT), urea, dodecyl-β-D-maltosideprotein (DDM), sodium chloride (NaCl), phosphatase inhibitor cocktail, standards including bovine serum albumin (BSA), β-casein were ordered from Sigma (St. Louis, MO). Titanium sulfate $Ti(SO_4)_2$ came from Sinopharm (Shanghai, China). Phosphoric acid were ordered from Kermel (Tianjin, China). Trifluoroacetic acid (TFA) was purchased from J&K Scientific (Beijing, China). NP-40 lysis buffer was ordered from Beyotime (Shanghai, China). Titansphere Phos-Tio Kit ($TiO_2$) was obtained from GL Sciences (Tokyo, Japan). Ni-NTA silica resin was purchased from (QIAGEN). Fourier-transform infrared (FT-IR) analysis was performed on an Tensor II FTIR spectrophotometer (Bruker, Germany). The surface morphologies were measured by a 400 field emission scanning electron microscope (SEM) (Bruker, Germany). Energy-dispersive spectroscopy (EDS) elemental mapping was performed using the Energy dispersive X-ray microanalysis system (Bruker, Germany).

**Cell and Oocyte Culture**

For studying interacting proteins of MAPK3, we constructed HEK-293A cell lines stably expressing pLVX-MAPK3-GFP-biotin (experiment group) and pLVX-GFP-biotin (control group) proteins. HEK-293A cells and HeLa cell were cultured in Dulbecco's modified Eagle's medium (DMEM) supplemented with 10% fetal bovine serum, 1% penicillin, and streptomycin in a humidified atmosphere of 5% $CO_2$ in air at 37 °C.

Wild-type C57BL/6 mice were purchased from the China Hunan Slac Jingda Laboratory Animal Co., Ltd. The mice were housed under specific pathogen-free (SPF) conditions in a controlled environment maintained at 20-22 °C, with a 12/12 hour light/dark cycle, 50-70% relative humidity, and provided with food and water ad libitum. To induce superovulation, 4-week-old female mice received an intraperitoneal injection of 5 IU pregnant mare serum gonadotropin (PMSG; Hangzhou Animal Pharmaceutical Co., Ltd.), followed 44 hrs later by an intraperitoneal injection of 5 IU human chorionic gonadotropin (hCG; Hangzhou Animal Pharmaceutical Co., Ltd.). Mouse MII oocytes were collected from oviducts 16 hours post-hCG administration. Oocytes were released into pre-warmed (37 °C) M2 medium (Sigma-Aldrich, cat# M7167) and subsequently cultured under mineral oil in microdrops of M16 medium at 37 °C in a humidified atmosphere of 5% $CO_2$. All procedures in this study were conducted in accordance with the Guide for the Care and Use of Laboratory Animals and were approved by the Animal Welfare Ethics Committee of Central South University (approval no. 2021sydw0263).

**Sample Preparation of Standard proteins.**

β-casein and BSA were dissolved in 8 M urea and 50 mM TEAB (pH 8.5) buffer. Adding 50 mM DTT solution to a final concentration of 2 mM and reduced at 37 °C for 30 min. For protein alkylation, a 100 mM IAA solution was added to reach a final concentration of 5 mM followed by incubating in the dark at room temperature for 30 min. The urea concentration in the solution was diluted 10-fold using 50 mM TEAB buffer (pH 8.5). Protein digestion was conducted with trypsin at a enzyme-to-substrate ratio of 1:50 (w/w) and incubated at 37 °C for 4 h. Subsequently, the same amount of trypsin was added again at 37 °C overnight. The digests were desalted with a C18 spin column, vacuum-dried in a SpeedVac (Thermo Scientific) and stored at -80 °C.

**Affinity Purification.**

We adopted the lentivirus system with character of puromycin resistance for constructing biotin tagged MAPK3 and GFP stable cell lines. The lentivirus system contains two plasmids: MAPK3 fused with GFP protein and biotin-binding protein at the C-terminus(pLVX-MAPK3-GFP-biotin) and pLVX-GFP-biotin. The lentivirus was generated in HEK293FT cell line for 48 hours after transfection of the two plasmids. The viruses were harvested by transferring the supernatants into a syringe packed with a 0.45μm filter. The 293A cell lines were infected with the viruses for 48 hours and subsequently treated with 3μg/mL of puromycin for the stable cell line selection.

Cells were lysed by IP lysis buffer (NP-40, 1% phosphatase inhibitor) and sonicated at 20 W for 5 s on/10 s off pulse until the solution became colorless. The solution was then centrifuged at 16,000 g for 20 min at 4 °C, with the supernatant collected for further processing. For protein enrichemnt, streptavidin sepharose were washed three times with 1 mL PBS and centrifugated (2000 g, 2 min). The sample was incubated with pre-washed streptavidin sepharose at 4 °C end over end ovenight. Next, the streptavidin sepharose were washed twice with NP-40 buffer and twice with 50 mM TEAB (pH 8.5) sequentially. Then, the streptavidin sepharose were heated at 95 °C for 5 min in 50 mM TEAB (pH 8.5). After cooling to room temperature, the sample was subjected to reduction (20 mM DTT) for 30 min at 65 °C and alkylation (40 mM IAA) for 20 min at room temperature in the dark. The sample was diluted to 1 mL with 50 mM TEAB (pH 8.5). Then, on-beads digestion was performed by adding trypsin and Lys-C overnight at 37 °C at an enzyme-to-protein ratio of 1:50 (w/w). After digesting, the streptavidin sepharose were washed by 200 μL of 50 mM TEAB (pH 8.5) and centrifued to get supernatant. The supernatant in two steps were merged and desalted using homemade C18 micro spin colume followed by drying down.

**Enrichment of Phosphopeptides by $TiO_2$ and Fe-NTA.**

The $TiO_2$ enrichment was performed according to the conditions described by Muneer et al.[1] with minor modifications. An equivalent volume of loading buffer A (3.2 M lactic acid, 60% ACN, 0.5% TFA) was added to the digest. Next, $TiO_2$ beads suspended in buffer B (80% ACN, 0.1% TFA) at a concentration of 75 μg/μL were added to the sample and incubated with shaking (1,500 rpm) at 25°C for 5 minutes. The mixture was centrifuged at 1,000g for 30 seconds to pellet the TiO2 beads, and the supernatant containing free peptides was discarded. The $TiO_2$ beads were then resuspended in 50 μL of wash buffer C (3.2 M lactic acid, 60% ACN, 0.1% TFA) and transferred to a C8 membrane D200-StageTip. An additional 50 μL of wash buffer C was added to the sample tube to collect any remaining beads, which were then transferred to the C8 membrane D200-StageTip. After centrifugation at 1,000 rpm for 1 minute to pass the solution through the tip, the beads were washed three times with buffer B. Finally, phosphopeptides were eluted with 50 μL of 50% ACN, 5% $NH_3 \cdot H_2O$. The eluted phosphopeptides were dried in a SpeedVac and stored at -80°C until LC-MS/MS analysis.

The Fe-NTA enrichment was performed according to the conditions described by Huang et al.[2] with minor modifications. Briefly, a homemade Fe-NTA enrichment tip column consists of one disk of C8 membrane and 10 μL of Fe-NTA Agarose. The tip was inserted into a 600 μL EP tube. The self-packed Fe-NTA tips were pre-equilibrated with 20 μL Fe-NTA loading/washing buffer (80% ACN, 0.1% TFA) by centrifugating at 200 g for 5 min for three times. Phosphopeptides in 80 μL loading buffer was evenly distributed and sequentially loaded to the Fe-NTA tips. Phosphopeptides were enriched by 600 rpm centrifugation at room temperature for 30 min twice. After being washed twice with Fe-NTA washing buffer, the columns were eluted with Fe-NTA elution buffer (60% ACN, 3% $NH_3 \cdot H_2O$). The eluted phosphopeptides were dried down with a SpeedVac and stored at -80 °C until LC-MS/MS analysis.

**MALDI-TOF Analysis**

Analysis of β-casein digest and MAQPFS(phospho)LR with or without interference of BSA digest was conducted by matrix-assisted laser desorption/ionization-time of flight (MALDI-TOF) MS on an Ultraflex III TOF/TOF mass spectrometer (Bruker Daltonics; Bremen, Germany). A matrix solution containing 2,5-dihydroxybenzoic acid (DHB, 25 mg/mL) was prepared in $ACN/H_2O/H_3PO_4$ (60:40:1, v/v/v). Equal volumes (2 μL) of the sample and DHB were sequentially deposited onto the MALDI plate for mass-spectrometric analysis. Spectra were acquired in positive-ionization mode using reflector detection.

**Data Analysis.**

MSFragger (version 22.0) and Proteome Discoverer 2.5 (Thermo Fisher Scientific) were used to search the DDA data. The FASTA database was downloaded from UniProt (Homo sapiens, TaxID 9606, 20468 proteins including common contaminants). The reversed proteins were used as decoys. FragPipe's "LFQ-phospho" workflow was used for the analysis. After filtering the results with global PSM- and protein-level 1% FDR, phosphosites with localization probability >0.75 were counted. Database searching of the raw files with Proteome Discoverer 2.5 was performed with the Sequest HT search engine. The database-searching parameters of phosphopeptides from HeLa cells lysates were set as below: full tryptic digestion and allowed up to two missed cleavages, the precursor mass tolerance was set at 10 ppm, whereas the fragment-mass tolerance was set at 0.02 Da. Carbamidomethylation of cysteines (+57.0215 Da) was set as a fixed modification, and variable modifications of methionine oxidation (+15.9949 Da), acetyl (N-terminus, +42.011 Da), and phosphorylation (serine (S), threonine (T) or tyrosine (Y), +79.966 Da) were allowed. The false-discovery rate (FDR) was less than 1%. The phosphosites with greater than 0.75 localization probabilities were considered as confident identification.

DIA Raw data were processed using DIA-NN software (version 1.9.1) for spectral library-free search and label-free quantitative analysis. The UniProt Mus musculus database (downloaded August 23, 2022; 55,319 entries) was utilized with the following parameters: Serine, Threonine or Tyrosine phosphorylation, Methionine oxidation, Protein N-terminal acetylation and methionine excision as variable modifications, carbamidomethylation of cysteine as a fixed modification, tryptic digestion allowing up to two missed cleavage, and peptide length restricted to 7-30 amino acids. A false discovery rate (FDR) threshold of ≤ 1% was applied, while other parameters in DIA-NN were maintained at default settings.

The kinase and phosphatase analyses were performed using the KinMap database (http://www.kinhub.org/kinmap/) and the human dephosphorylation database (DEPOD) (www.depod.org), respectively. The kinase families listed includes TK (tyrosine kinases), TKL (tyrosine kinase-like), CK1 (casein kinase 1), CAMK (calcium/calmodulin-dependent protein kinase), AGC (containing PKA, PKG, PKC families), CMGC (containing CDKs, MAPK, GSK, CLK families), and STE (serine/threonine kinases many involved in MAPK kinases cascade). The phosphatase family abbreviated are R-PTP (receptor protein tyrosine phosphatases), NR-PTP (nonreceptortype protein tyrosine phosphatases), DSP (dual specificity phosphatase), EYA (eyes absent), HAP (histidine acid phosphatase), PPM (protein phosphatase $Mg^{2+}$ or $Mn^{2+}$ dependent), FCP/SCP (TFIIF-associating component of RNA polymerase II CTD phosphatase/small CTD phosphatase), PM (phosphoglycerate mutase), dNT (5'(3')-deoxyribonucleotidase), CDC25 (cell division cycle 25) and PGAM (Phosphoglycerate mutase), Lipin (Lipin Phosphatidate), PHLP (pH domain leucine-rich repeat protein) and PPP (phosphoprotein phosphatase).

Kinase enrichment analysis was performed using PhosphoSitePlus (v6.8.2) [3] to predict upstream kinases whose activity may underlie the phosphorylation changes during oocyte maturation. Phosphorylation sites showing significant upregulation in three comparative stages (GVBD vs. GV, MI vs. GV, and MII vs. GV) were analyzed against non-altered phosphosites via Fisher's exact test. The enrichment results are presented in a heatmap where color intensity indicates the enrichment value and asterisks denote statistical significance: ***$p < 0.001$, **$p < 0.01$, *$p < 0.05$. The associated kinome tree was visualized and annotated using KinMap [4].

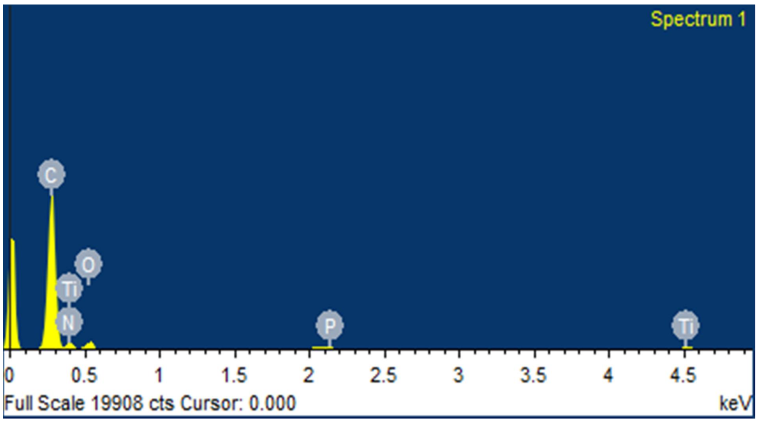


| Element | Weight% | Atomic% |
|---|---|---|
| C K | 69.51 | 75.14 |
| N K | 12.44 | 11.53 |
| O K | 15.59 | 12.65 |
| P K | 0.11 | 0.05 |
| Ti K | 2.35 | 0.64 |
| Totals | 100.00 | |

**Figure S1.** EDX spectrum and chemical composition (table) of Ti-PAN material.

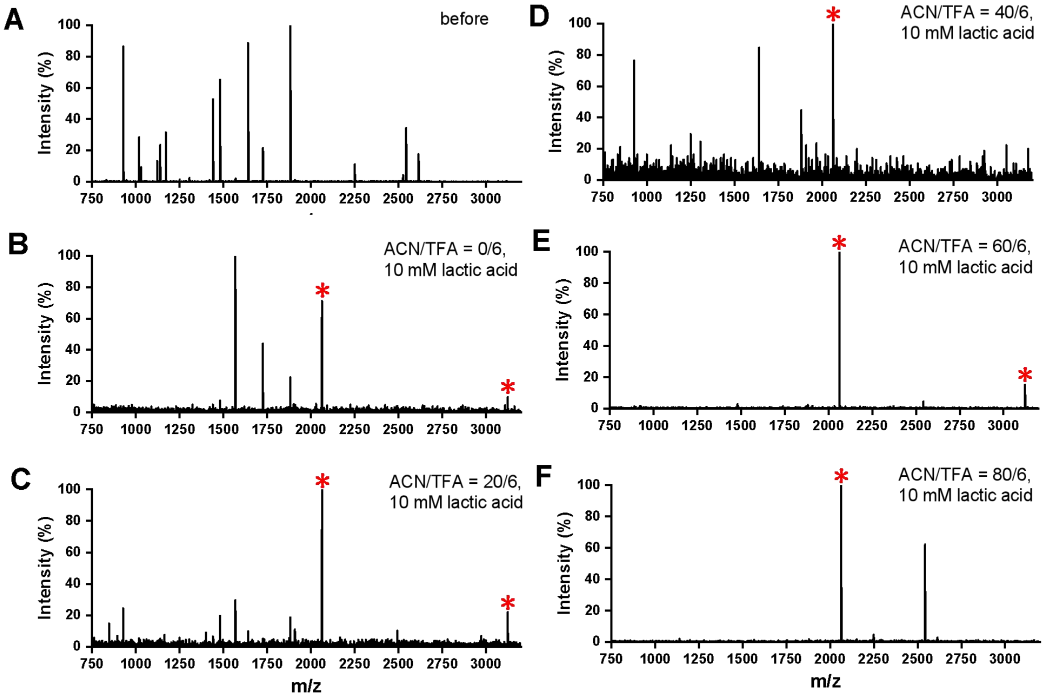


**Figure S2.** MALDI-TOF mass spectra of β-casein and BSA tryptic digest mixtures (A) before and (B-F) after enrichment by Ti-PAN with different loading buffer solutions: 0.6% TFA, 0.01 M lactic acid and (B) 0%, (C) 20%, (D) 40%, (E) 60%, (F) 80% ACN. Molar ratio of β-casein to BSA is 1:100. "*" indicates phosphopeptide.

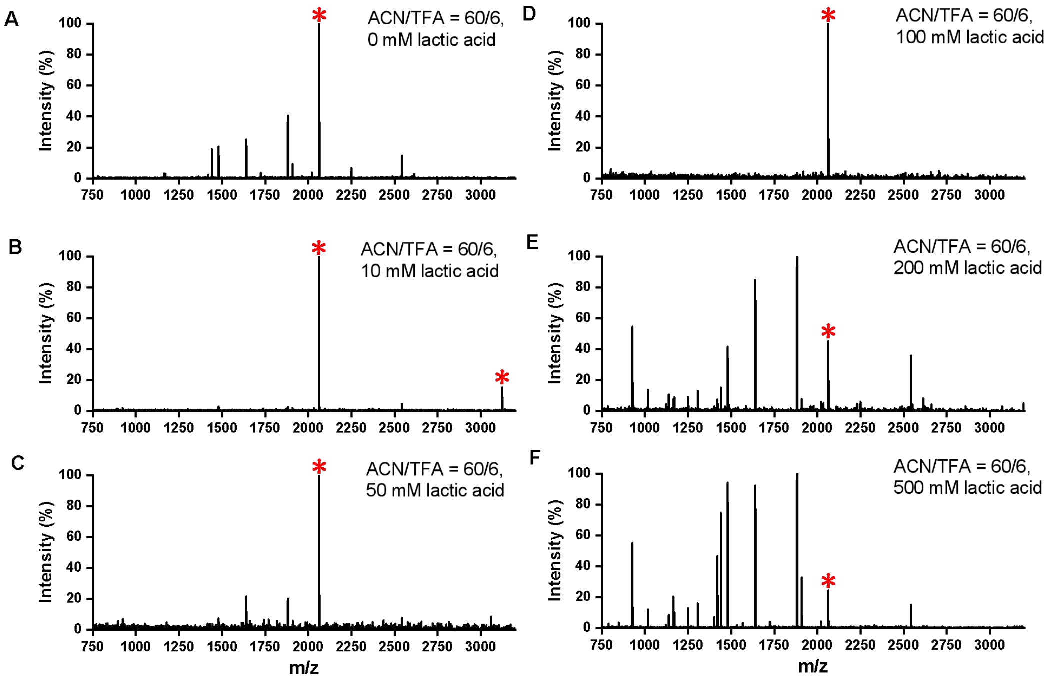


**Figure S3.** MALDI-TOF mass spectra of β-casein and BSA tryptic digest mixtures after enrichment by Ti-PAN with different loading buffer solutions: 60%ACN, 0.6% TFA and (A) 0 mM , (B) 10 mM, (C) 50 mM, (D) 100 mM, (E) 200 mM, (F) 500 mM lactic acid. Molar ratio of β-casein to BSA is 1:100. "*" indicates phosphopeptide. The optimal concentration of lactic acid in the loading buffer for phosphorylated peptide enrichment is 10 mM. Subsequently, with the increase of lactic acid concentration, the non-specific adsorption of Ti PAN was enhanced, resulting in a decrease in the enrichment ability of Ti PAN for phosphorylated peptides.

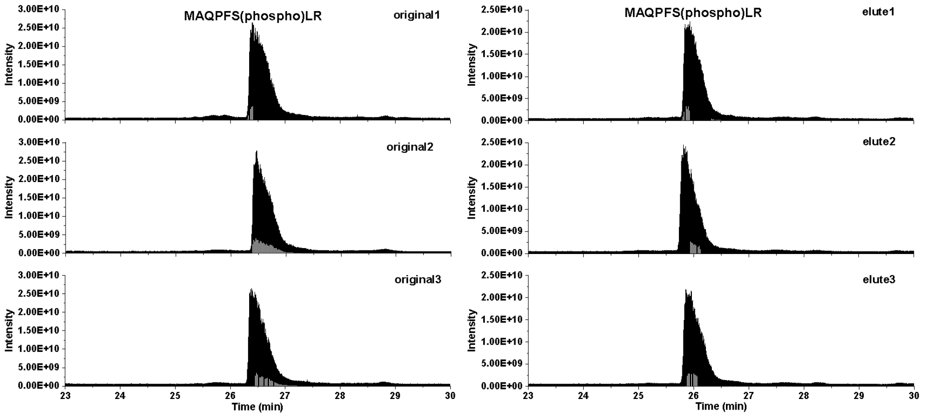


| | Apex RT | Start RT | End RT | Area | Height |
|---|---|---|---|---|---|
| elute1 | 25.94 | 24.8 | 28.42 | 2.82082E+11 | 22036498960 |
| elute2 | 25.81 | 24.93 | 28.6 | 3.14788E+11 | 24506550379 |
| elute3 | 25.85 | 24.88 | 28.38 | 3.12558E+11 | 21915501111 |
| origi1 | 26.4 | 26.14 | 29.28 | 3.52451E+11 | 26435515677 |
| origi2 | 26.48 | 26.33 | 28.7 | 3.39618E+11 | 27746654020 |
| origi3 | 26.39 | 26.14 | 28.57 | 2.83081E+11 | 25784555010 |

**Figure S4.** Evaluation of the phosphopeptide (MAQPFpSLR) recovery on the Ti-PAN tip.

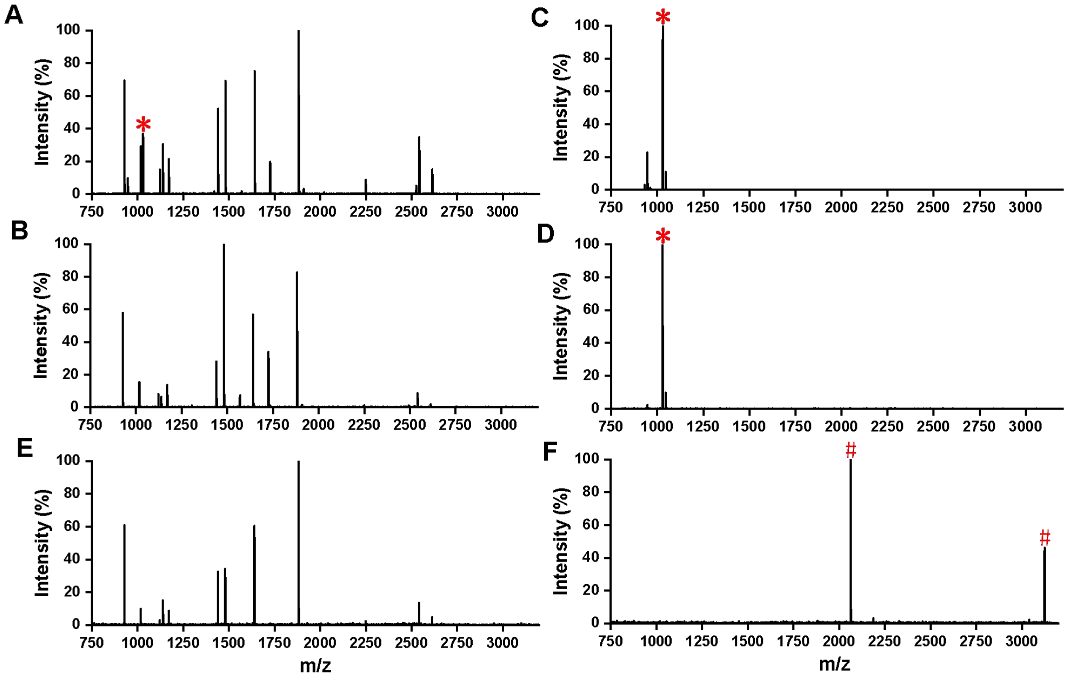


**Figure S5.** MALDI-TOF MS spectra of the mixture of BSA and (A-D) standard peptide or (E, F) β-casein at different molar ratios. Direct analysis at (A, E) 10:1 and (B) 100:1 ratio. Postenrichment at ratios of (C, F) 10:1 and (D) 100:1 ratio. Phosphopeptides are marked with red "*" of tryptic digests from standard peptide and red "#" of β-casein. As shown in figures of A, B, E, prior to enrichment, the signals of phosphorylated peptides were almost obscured due to their low abundance. After phosphopeptide enrichment, the signal intensity of phosphorylated peptides was significantly enhanced.

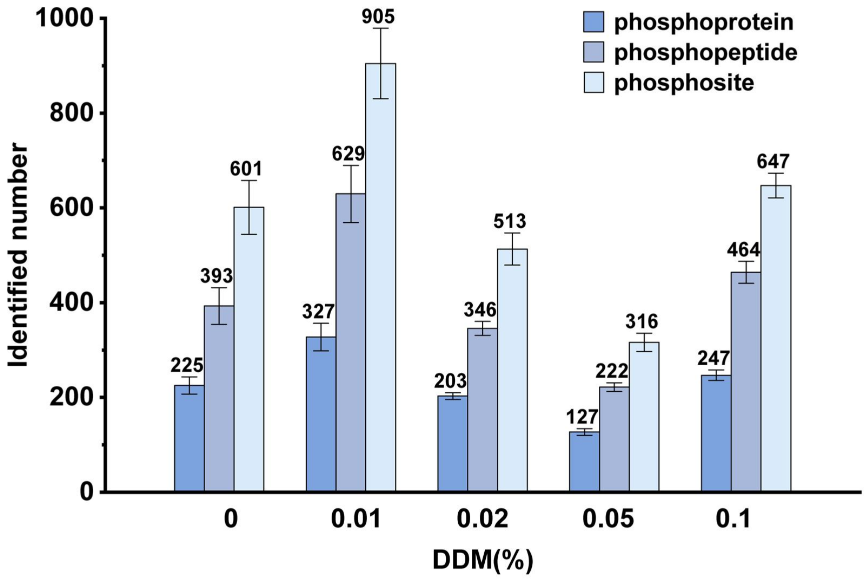


**Figure S6.** The number of identified phosphoproteins, phosphopeptides, and phosphosites at different concentrations of DDM.

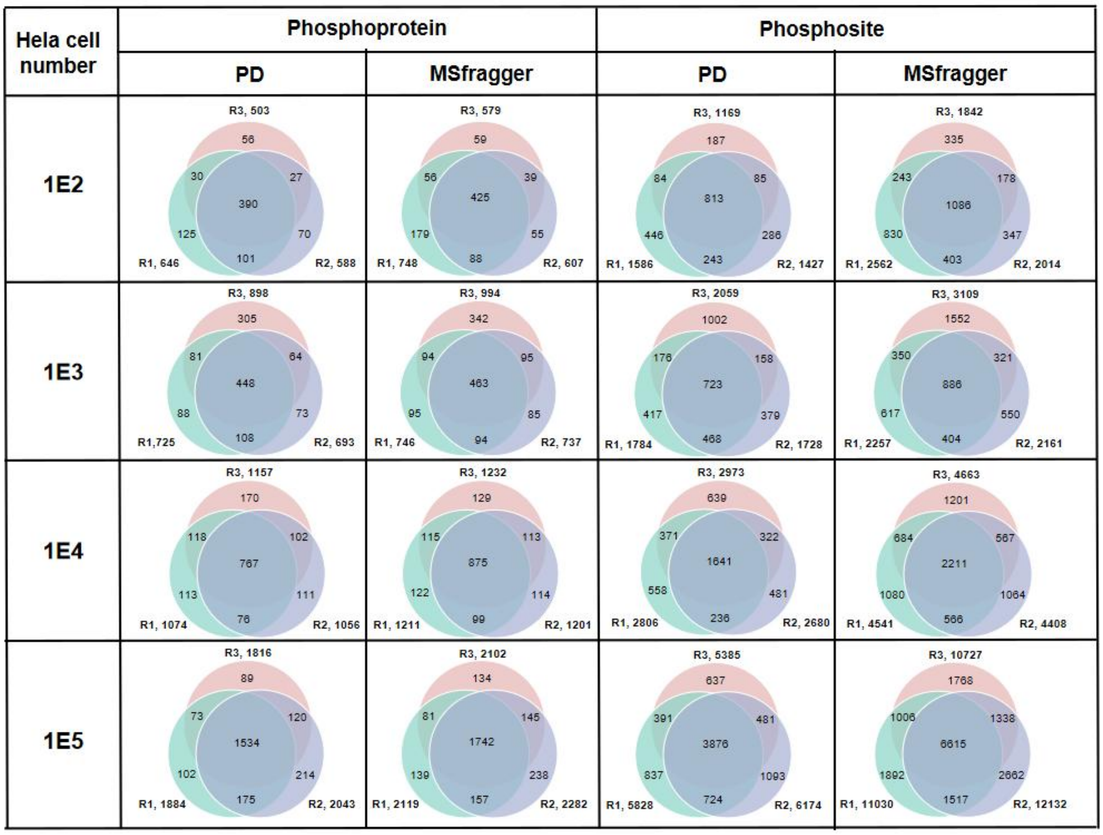


**Figure S7.** The Venn plots of identified phosphoproteins and phosphopeptides number from 1E2 to 1E5 Hela cells via Proteome Discoverer and MSFragger.

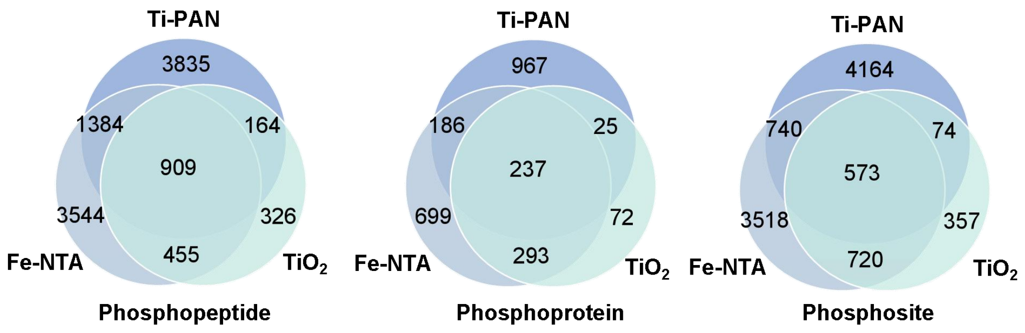


**Figure S8.** The Venn plots of identified phosphoproteins, phosphosites and phosphopeptides from 10,000 Hela cells with different materials via MSFragger.

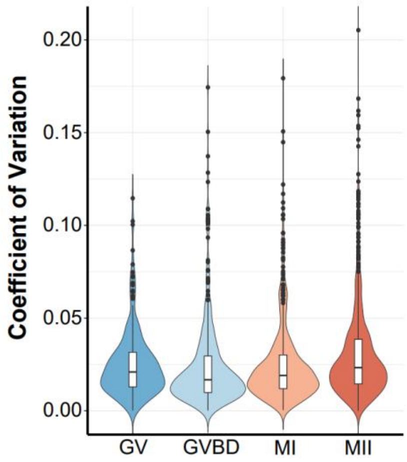


**Figure S9.** Coefficient of variation distribution of the LFQ intensity from five replicates at differnt stages.

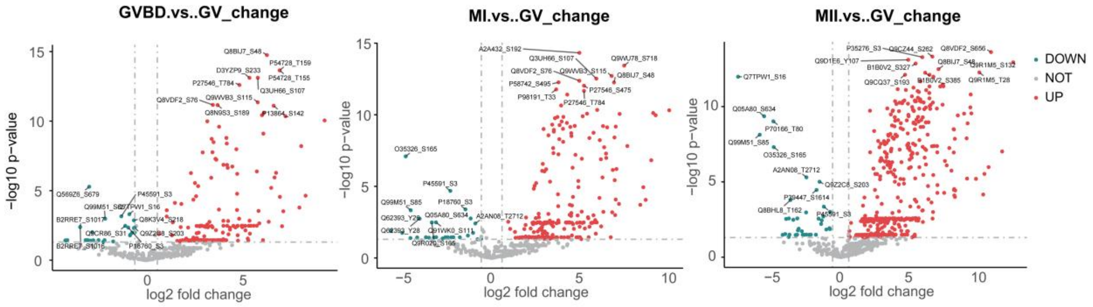


**Figure S10.** Volcano plots showing the differentially phosphorylated proteins between GVBD and GV (left), or between MI and GV (middle), or between MII and GV (right).

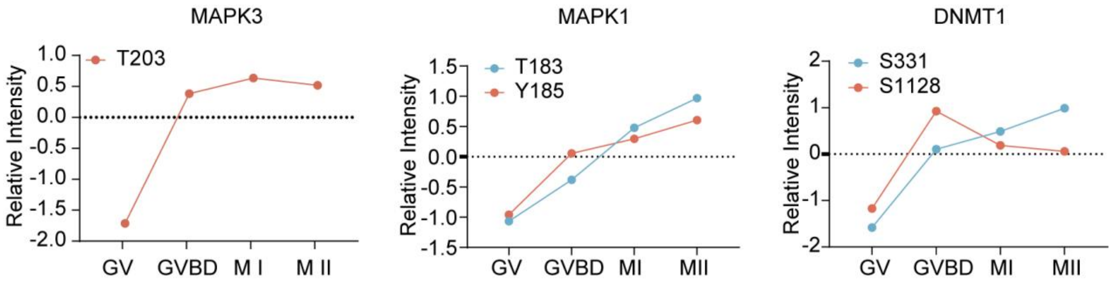


**Figure S11.** Relative abundance of phosphosites in representative proteins at different stages, with red and blue colors denoting sites newly identified in this study and those not previously detected in mouse oocytes, respectively.
